# Cryo-EM structures of SERCA2b reveal the mechanism of regulation by the luminal extension tail

**DOI:** 10.1101/2020.03.28.012849

**Authors:** Yuxia Zhang, Michio Inoue, Akihisa Tsutsumi, Satoshi Watanabe, Tomohiro Nishizawa, Kazuhiro Nagata, Masahide Kikkawa, Kenji Inaba

**Affiliations:** Institute of Multidisciplinary Research for Advanced Materials, Tohoku University, Sendai 980-8577, Japan; Graduate School of Medicine, The University of Tokyo, 7-3-1 Hongo, Bunkyo-ku, Tokyo 113-0033, Japan; Graduate School of Science, The University of Tokyo, 7-3-1 Hongo, Bunkyo-ku, Tokyo113-0033, Japan; Faculty of Life Sciences, Kyoto Sangyo University, Kyoto 603-8555, Japan; Core Research for Evolutional Science and Technology (CREST), Kawaguchi, Japan

## Abstract

SERCA2b is a Ca^2+^-ATPase that pumps Ca^2+^ from the cytosol into the ER and maintains the cellular calcium homeostasis. Herein, we present cryo-EM structures of human SERCA2b in E1·2Ca^2+^-AMPPCP and E2-BeF_3_^-^ states at 2.9 and 2.8 Å resolutions, respectively. The structures revealed that the luminal extension tail (LE) characteristic of SERCA2b runs parallel to the lipid-water boundary near the luminal ends of transmembrane (TM) helices TM10 and TM7, and approaches the luminal loop flanked by TM7 and TM8. Upon deletion of the LE, the cytosolic- and TM-domain arrangement of SERCA2b resembled that of SERCA1a, resulting in multiple conformations. The LE regulates the conformational transition between the E1·2Ca^2+^-ATP and E2P states, explaining the different kinetic properties of SERCA2b from other isoforms lacking the LE.

**One sentence summary:** Cryo-EM structures of SERCA2b at 2.8-2.9 Å resolutions reveal how the luminal extension tail regulates its activity.

---

The endoplasmic reticulum (ER) is the organelle where secretory and membrane proteins are synthesized and acquire higher-order structure under the surveillance of cellular protein quality control systems (*1, 2*). Another important function of the ER is calcium ion (Ca^2+^) storage, and regulation of Ca^2+^ efflux from this organelle plays a critical role in controlling cellular growth, proliferation, differentiation, and death (*3*). Thus, the ER has developed the elaborate mechanisms for maintaining Ca^2+^ homeostasis that utilize multiple kinds of Ca^2+^ pumps and channels.

Sarco/endoplasmic reticulum calcium ion ATPase (SERCA) is a P-type ATPase family member that conducts Ca^2+^ uptake from the cytosol to the ER against an uphill gradient of >1,000-fold difference in Ca^2+^ concentration (4). The SERCA family contains three isoforms SERCA1-3, among which SERCA2b serves as a ubiquitously expressed housekeeping enzyme involved in the cellular Ca^2+^ homeostasis. They share similar overall structure with three cytoplasmic domains comprising an actuator (A), a nucleotide-binding (N), and a phosphorylation (P) domain, and 10 transmembrane (TM) helices (Fig. 1A) that can be divided into three TM helix clusters, TM1-TM2, TM3-TM4, and TM5-TM10 (5, 6).

**Fig. 1.**
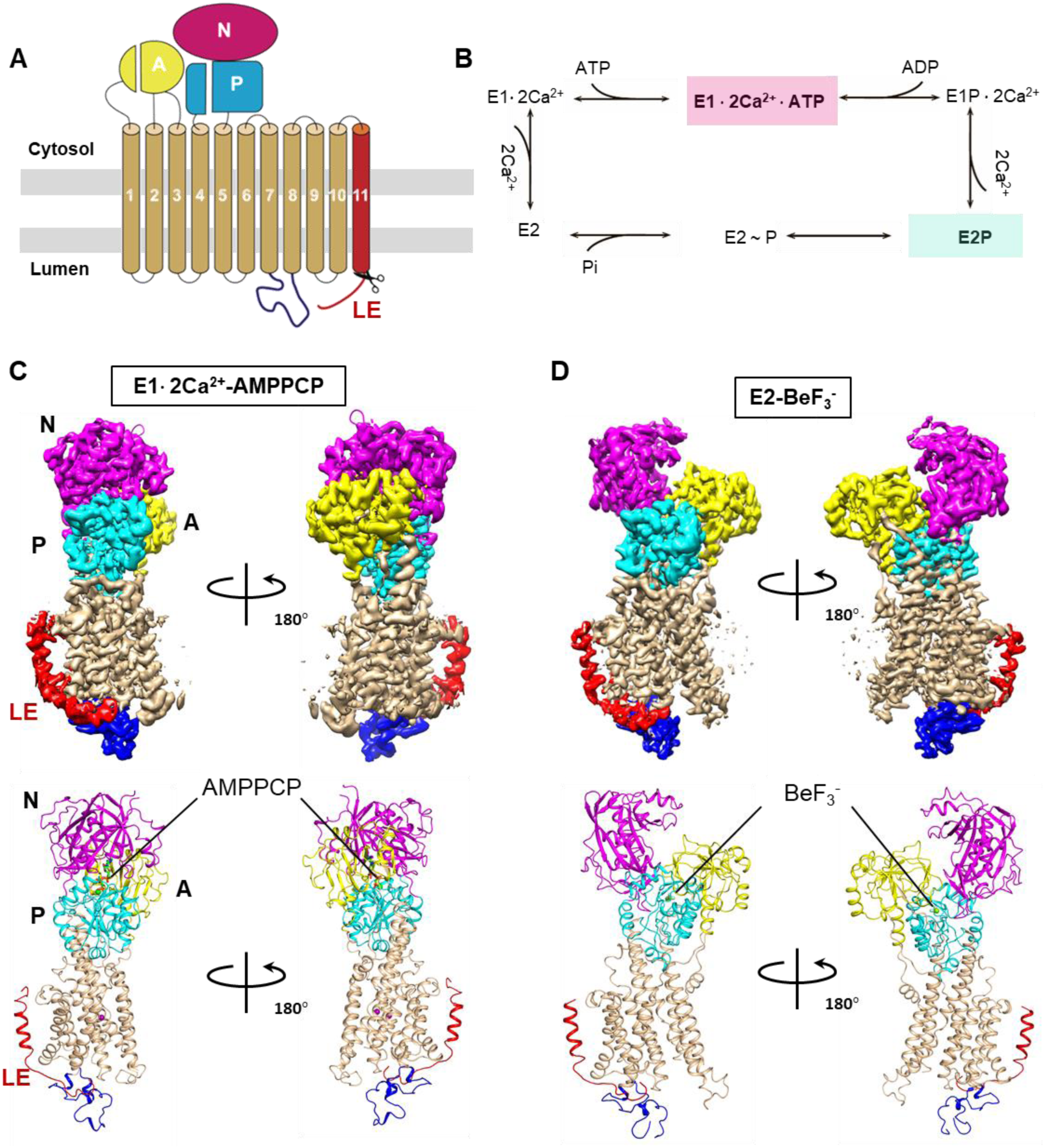
Cryo-EM structures of SERCA2b WT in E1·2Ca^2+^-ATP and E2P states. **A**. Topology diagram of SERCA2b. Conserved A, N, and P domains and transmembrane helices (TM1-TM10) are colored yellow, magenta, cyan and wheat, respectively. The C-terminal 11^th^ TM helix (TM11) and the subsequent luminal extension tail (LE) are colored red while L7/8 is colored blue. The scissors indicate the truncation site used to generate the T1032stop construct. **B**. Catalytic cycle for SERCA to transport Ca^2+^ from the cytosol to the ER lumen through hydrolysis of ATP. The intermediate sates for which cryo-EM structures have been determined in this work are colored pink and cyan, respectively. **C, D**. Overall cryo-EM structures of SERCA2b WT in E1·2Ca^2+^-AMPPCP (C) and E2-BeF_3_^-^ (D) states. The color pattern is the same as in A.

Crystal structures of a series of reaction intermediates and transition states stabilized with appropriate ligands or analogs (*7, 8*) have demonstrated the mechanism of actions of SERCA1a, an isoform specifically expressed in skeletal muscle, thereby revealing molecular details of the catalytic cycle of this Ca^2+^ pump. By contrast, structural and mechanistic information of SERCA2b remain scarce despite its physiological importance. SERCA2b shares 85% amino acid sequence similarity with SERCA1a, and shares residues 1-993 with SERCA2a, a splicing variant of SERCA2b. Unlike other isoforms, SERCA2b has a 49-residue C-terminal extension comprising a cytosolic loop (L10/11; Gly994-Asp1012), an additional (11^th^) TM helix (TM11; Gly1013-Tyr1030) and a luminal extension tail (LE; Ser1031-Ser1042). In this connection, SERCA2b exhibits a significantly lower maximal turnover rate than SERCA1a and SERCA2a, and almost 2-fold higher apparent calcium affinity (*9*).

Our recent crystal structure of SERCA2b at 3.45 Å resolution revealed that, despite discrepancy with a previously reported structural model (*10*), TM11 is located adjacent to TM10, and engages in weak interactions with a part of the L8/9 loop and the N-terminal end of TM10 (*11*). Structural comparisons further demonstrated that due to the presence of TM11, SERCA2b has a different TM helix orientation relative to the cytosolic domains from those of SERCA2a and SERCA1a. However, electron density was missing for several critical segments including the LE. Additionally, crystal structure of SERCA2b was solved only for the E1-2Ca^2+^-adenylyl methylenediphosphonate (AMPPCP)-bound state. Thus, the mechanism underlying the SERCA2b regulation by the C-terminal extension remains poorly understood.

In the present work, we determined cryo-EM structures of human SERCA2b in E1·2Ca^2+^-AMPPCP and E2-BeF_3_^-^ states at resolutions of 2.9 Å and 2.8 Å, respectively. The higher-resolution structures illuminated both the backbone and side-chain conformations over almost the entire part of SERCA2b. Furthermore, to define the location of the LE and gain deep insight into its regulatory roles, we determined cryo-EM structures of SERCA2b T1032stop, a truncated construct that spans the protein chain up to TM11 but lacks the LE. Structural comparison between wild-type (WT) SERCA2b and the T1032stop variant reveals the exact location of the LE and the mechanism of the LE-mediated structural regulation in SERCA2b. The present findings explain the regulated conformational transition between the E1·2Ca^2+^-ATP and E2P states in SERCA2b.

## Overall structures of SERCA2b in E1**·**2Ca^2+^-AMPPCP and E2-BeF_3_^-^ states

We expressed and purified SERCA2b WT and T1032stop essentially as described previously (*11*) (Fig. S1), except that the buffer used for the final exchange comprised 50 mM HEPES (pH 7.0), 100 mM KCl, and 0.01% (w/v) lauryl maltose neopentyl glycol (LMNG), but contained no glycerol to allow acquisition of high-quality particle images. The cryo-EM structures of SERCA2b WT in E1·2Ca^2+^-AMPPCP and E2-BeF_3_^-^ states were determined as mimics of the E1·2Ca^2+^-ATP and E2·P states (Fig. 1B) at a resolution of 2.9 Å and 2.8 Å, respectively (Table S1).

In the cryo-EM structure of SERCA2b WT in the E1·2Ca^2+^-AMPPCP state, all cytosolic domains and TM helices including TM11 are located at almost the same positions as observed in the previous crystal structure (*11*) (Fig. 1C and Fig. S2A). The overall RMSD of Cα atoms between these two structures is 1.10 Å. The higher-resolution cryo-EM structures provided more precise views of the modes of AMPPCP and Ca^2+^ binding (Fig. S3A and B). Moreover, no-alignment 3D classification with a mask covering TM11 resulted in clear density for the entire part of TM11 and its neighboring regions, even though density was still invisible for the major portion of L10/11 probably due to its intrinsically disordered structure. Thus, the cryo-EM structures of SERCA2b verified the close contacts between the side chains of W927 (L8/9) and Phe1018 (TM11) and between those of L967/L970/K971 (TM10) and V1029 (TM11) (Fig. S2B), of which regulatory roles were proposed previously (*11*).

Notably, the LE of SERCA2b was visible in the cryo-EM maps of both the E1·2Ca^2+^-AMPPCP and E2-BeF_3_^-^ states (Fig. 1C, D and Fig. S4A and B). In sharp contrast, SERCA2b T1032stop did not show density around the corresponding regions (Fig S4C and D), corroborating our identification of the LE. Consequently, the LE has been found to locate near the luminal ends of TM10 and TM7, and approach the short α-helix (Phe866-Ser870) in L7/8 (Fig. 2A and B). The actual location of the LE strikingly differs from the putative site in the previous docking model, in which the loop was predicted to reside in the groove between L5/6 and L7/8 (*10*).

**Fig. 2.**
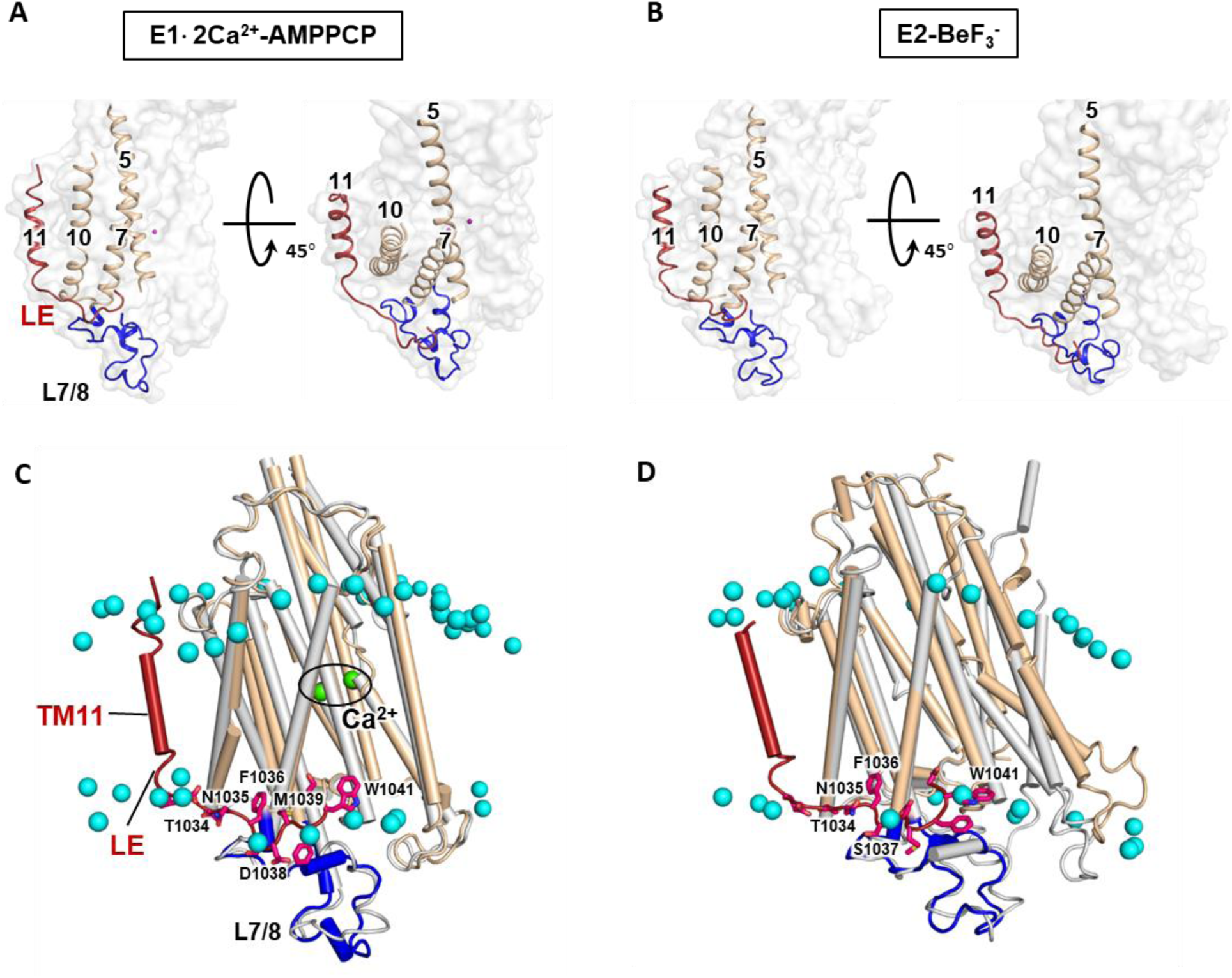
Location of the LE in SERCA2b. **A, B**. Location of the LE in the E1·2Ca^2+^-AMPPCP (A) and E2-BeF_3_^-^ (B) states. The TM11-LE and L7/8 segments are colored red and blue, respectively. **C, D.** Location of the LE relative to the first layer of phospholipids identified in crystal structures of SERCA1a. The cryo-EM structures of SERCA2b are superimposed to crystal structures of SERCA1a with the surrounding phospholipids in the E1 · 2Ca^2+^-AMPPCP (C) and E2-BeF_3_^-^ (D) states (PDB ID: 5XA8 and 5XA9, respectively) such that the RMSD of Cα atoms in TM7-TM10 is minimized. Phosphorous atoms of the phospholipids are represented with cyan spheres. The color pattern for SERCA2b is the same as in panel A, while the crystal structures of SERCA1a are colored gray.

In the preceding work (*12*), the entire first layer of phospholipids surrounding the transmembrane helices of SERCA1a was visualized in the electron density maps obtained by X-ray solvent contrast modulation. By superimposing the present cryo-EM structures of SERCA2b to the crystal structures of SERCA1a with the surrounding phospholipids, we found that the LE runs parallel to the lipid-water boundary in the luminal side (Fig. 2C &D). Accordingly, the LE contains polar and charged residues on the aqueous side, whereas the hydrophobic residues including F1036 and W1041 are directed toward the interior of the lipid bilayer. The LE contains no basic amino acids, which could, if any, anchor the nearby phospholipids through salt bridges with phosphate groups. It therefore seems unlikely that the LE serves to fix particular phospholipids or locally distort the lipid bilayer in accordance with the TM helix movements during the SERCA reaction cycle. Rather, the LE likely acts as an element that directly regulates the location and orientation of both the cytosolic and transmembrane domains of SERCA2b, as described later.

## Conformational transition from E1**·**2Ca^2+^-ATP to E2P states in SERCA2b

The previous crystallographic studies of SERCA1a demonstrated that during the transition from E1·2Ca^2+^-ATP to E2P states, the nucleotide-binding site in the N-domain and the Ca^2+^ exit channel constituted by TM4-6 are opened to the cytosolic and luminal sides, respectively, to facilitate the release of ADP and Ca^2+^ (*5, 13*). The present study revealed that, similarly to SERCA1a, AMPPCP binds to the boundary of the P- and N-domains of SERCA2b, resulting in a compact headpiece cluster composed of the cytosolic A-, N- and P-domains (Fig. 1C and Fig. 3). In the E2-BeF_3_^-^ state, only a BeF_3_^-^ molecule is covalently linked to the carboxylate side chain of Asp351 with assist of a nearby Mg^2+^ ion (Fig. S3C). Due to the dissociation of the ADP moiety located at the inter-domain interface, the cytoplasmic domains in the E2-BeF_3_^-^ state are more separated from each other than those in the AMPPCP-bound state. Thus, the A-, N- and P-domains in the E2-BeF_3_^-^ state are tilted down toward the membrane and rotated by 99°, 60° and 20°, respectively, relative to the corresponding domains in the E1·2Ca^2+^-AMPPCP state (Fig. 1D and Fig. 3). The movement of the A-domain imposes a force on TM1-2 in a lateral direction, which relocates TM3-4 through van der Waals contacts between TM1/TM3 and the luminal half of TM4 (Fig. 3, right lower inset). Eventually, TM1-4 are separated from TM5-6 to open the calcium exit channel, whereas the other TM helices (TM7-10) constitutes a more rigid cluster. Thus, SERCA2b undergoes a significant rearrangement in both the cytosolic and TM domains upon ATP hydrolysis and subsequent ADP dissociation, leading to the facilitated release of Ca^2+^, as previously observed for SERCA1a.

**Fig. 3.**
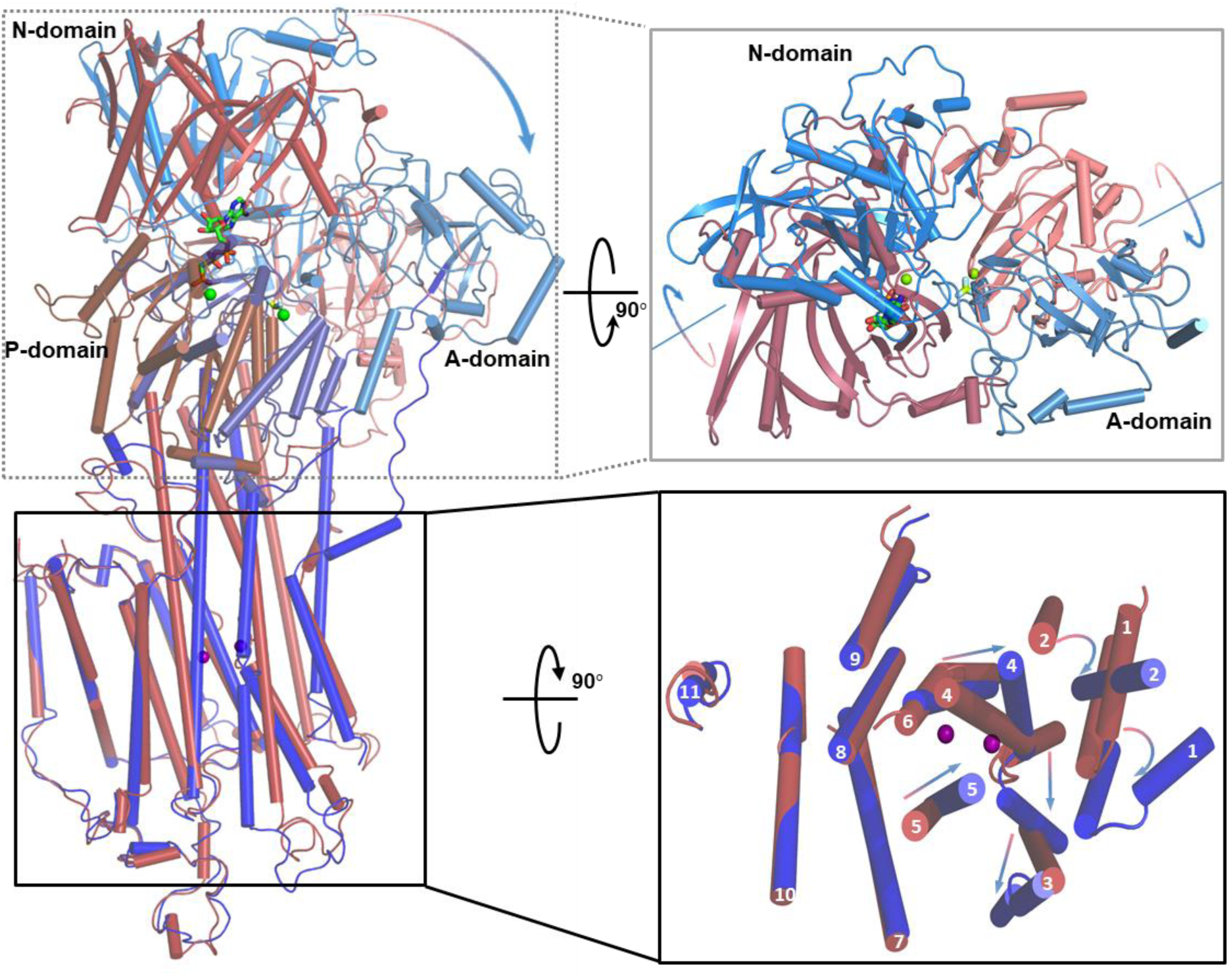
Conformational transition of SERCA2b from E1·2Ca^2+^-ATP to E2P states. Cryo-EM structures of SERCA2b WT in the E1 · 2Ca^2+^-ATP and E2 · P states are superimposed such that the RMSD of Cα atoms in TM7-TM10 is minimized. Helices are represented by cylinders. Right upper and lower panels show views of SERCA2b from cytosolic and luminal sides, respectively. A, N, P, and TM-helix domains in the E1·2Ca^2+^-AMPPCP state are colored salmon, crimson, brown, and red, respectively. Those in the E2-BeF_3_^-^ state are colored light-blue, marine, deep-blue and blue, respectively. Arrows indicate the rotation or movement of the cytosolic domains and TM helices during the transition from E1 · 2Ca^2+^-ATP to E2P states. AMPPCP and BeF_3_^-^ molecules are represented by green and cyan sticks, respectively. Bound magnesium and calcium ions are depicted as green and purple spheres, respectively.

## Roles of the LE in structural regulation of E1·2Ca^2+^-ATP form of SERCA2b

Detailed structural comparison of SERCA2b WT and T1032stop provided deep insight into the regulatory role of the LE. Whereas the cryo-EM structure of WT revealed a single conformation characteristic of SERCA2b, the T1032stop mutant was classified into three different classes of conformations in a similar abundance (class1 = 26.7 %, class2 = 26.7 %, class 3 = 23.5 %) after the additional local search 3D classification with masks to remove micelle density (Fig. 4A). The same data processing did not yield different classes of conformations for WT. All classes clearly showed density of bound AMPPCP, its neighboring Mg^2+^, and two bound Ca^2+^ in the cryo-EM maps (Fig. S5A and B), indicating that the preparation of E1·2Ca^2+^-AMPPCP form was homogeneous. Like the wild-type protein, T1032stop retained the compact headpiece cluster of the A-, N- and P-domains in all classes (Fig. 4A). However, these cytoplasmic domains stood more upright relative to the TM domain. Structural comparison between these three classes demonstrated that while L6/7 in class 1 approached more closely to L8/9 than that in the other two, the P-domain in class 1 was situated even more upright relative to the membrane (Fig. 4A, right inset), suggesting that the locations of L6/7 and L8/9 are linked to the orientations of the cytosolic domains.

**Fig. 4.**
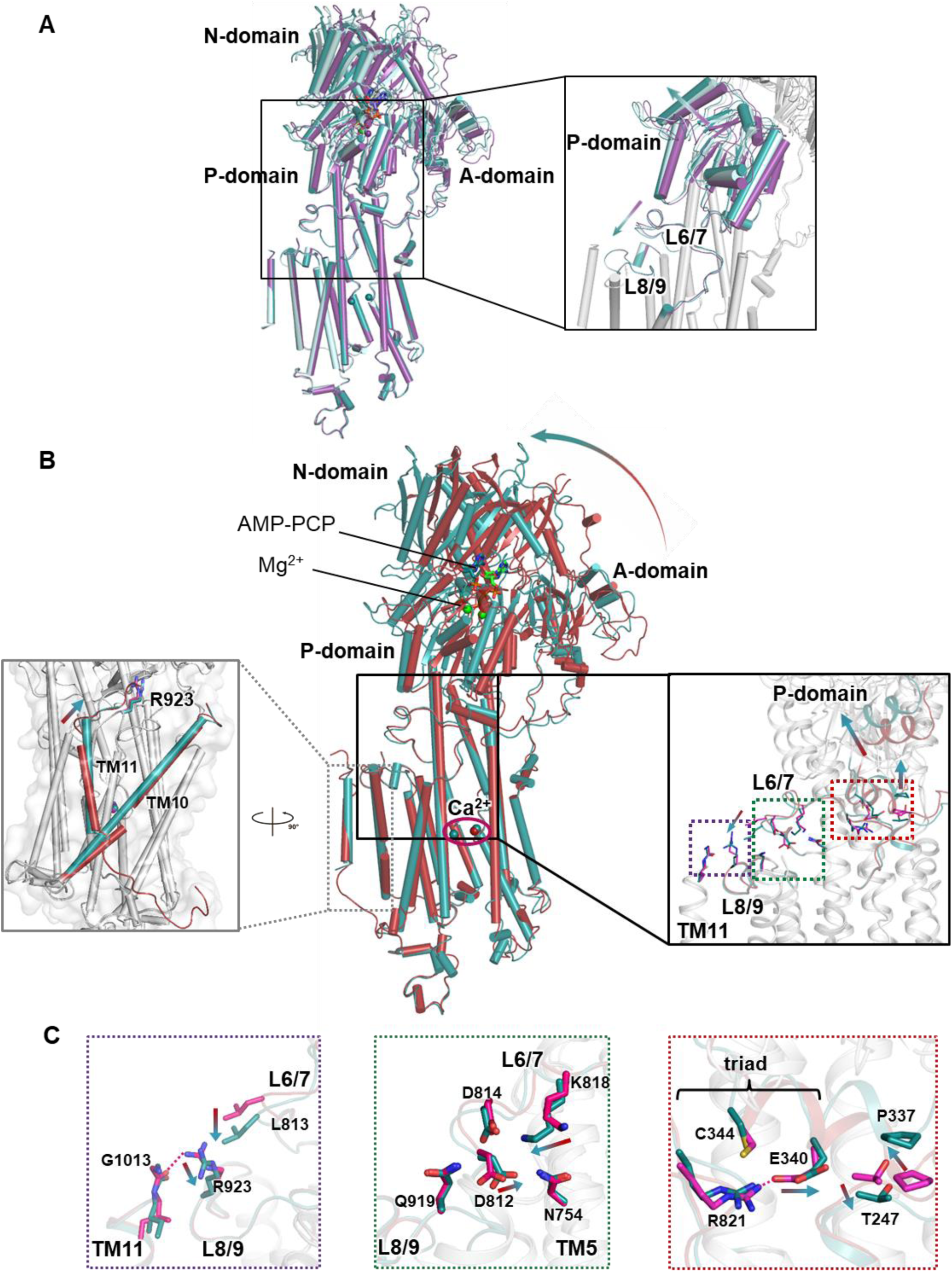
Structural effects of the LE on SERCA2b in the E1·2Ca^2+^-ATP state. **A**. Cryo-EM structures of three different classes (class 1, 2 and 3) of SERCA2b T1032stop in the E1·2Ca^2+^-AMPPCP state are superimposed such that the RMSD of Cα atoms in TM7-TM10 is minimized. Class 1, 2 and 3 are colored dark-cyan, pale-cyan, and violet, respectively. A right inset highlights the linkage of the L6/7 movement to the relocation of P-domain. **B.** Cryo-EM structures of SERCA2b WT and class 1 of T1032stop in the E1·2Ca^2+^-AMPPCP state are superimposed such that the RMSD of Cα atoms in TM7-TM10 is minimized. Cylinder diagrams of SERCA2b WT and class 1 of T1032stop are colored red and cyan, respectively. The right inset presents a close-up view of the cytosolic parts of TM6-11, L6/7, and L8/9. The left inset shows a side view of the TM-helix domain, in which TM10 and TM11 are indicated by red (WT) and cyan (class 1 of T1032stop) cylinders. Residues critical for the positional shifts of the cytosolic domains and TM helices upon deletion of the LE are represented by sticks in the right inset. The movements of the cytosolic domains and TM helices induced by deletion of the LE are indicated by arrows in both insets. **C.** The regions surrounded by violet, green and red dotted squares in the right inset of panel B are highlighted in the left, middle and right panels, respectively. The movements of residues in L6/7 and P-domain induced by deletion of the LE are indicated by arrows. Red dotted lines in the left and right panels indicate a hydrogen bond between the R923 side chain and the G1013 main-chain carbonyl group and a salt bridge between the side chains of E340 and R821 formed in SERCA2b WT.

In this context, deletion of the LE relocated L813 in L6/7 closer to R923 in L8/9 to make van der Waals contact between these two (Fig. 4C, left panel). The concomitant approach of L6/7 to L8/9 appears to influence the conformation of a triad constituted by R821 in L6/7, E340 and C344 in the P-domain (Fig. 4C, right panel), accompanying the slight but significant repositioning of the intermediary residues (N754, D812, D814, K818, and Q919) (Fig. 4C, middle). Whereas this triad is highly conserved in P-type ATPases (*14*) and supposed to link the P-domain to L6/7 and the cytoplasmic part of TM3 through hydrogen bonds (*15*), the side chain of E340 is significantly more apart from those of R821 and C344 in class 1 of T1032stop (Fig 3C, right and Fig. S6A). Additionally, P337 in the P-domain and T247 in TM3 are markedly repositioned (Fig 3C, right and Fig. S6A). These local conformational changes at the triad and its neighboring region allow the P-domain to move away from the original position (Fig. 4B, right inset), resulting in the upright rotation of the A- and N-domains (Fig. 4A) as the P-domain provides a flat surface for tight interactions with the A-domain, and the bound AMPPCP molecule serves as a bridge that forces the N-domain to move in accordance with the P-domain. The triad thus altered likely allowed generation of the three different classes in T1032stop.

In line with this notion, while SERCA1a retains a hydrogen bond between R822 (R821 in SERCA2b) and Cys344, its E340 residue is situated well far from these two residues (*15*) and renders the cytoplasmic domain cluster more upright with respect to the membrane as in T1032stop. As a result, the overall domain arrangement of SERCA1a is highly superimposable to that of class 1 of SERCA2b T1032stop (Fig. S7A). Thus, the overall conformation of SERCA2b resembled that of SERCA1a upon deletion of the LE. In support of this, the RMSD values for Cα atoms between class 1 of SERCA2b T1032stop and SERCA1a is 0.602 Å, significantly smaller than between class 1 of T1032stop and SERCA2b WT (1.678 Å). Meanwhile, the other classes (class 2 and class 3) obviously adopt overall conformations intermediate between those of SERCA2b WT and class 1 of T1032stop (Fig. 4A).

Deletion of the LE also altered the orientation of TM11 (Fig. 4B, left inset), which likely influenced the positions of the N-terminal end of TM11 and hence the locations of R923 and L6/7 (Fig. 3C, left), as described above. In this connection, the hydrogen bond formed between the R923 side chain and the G1013 main-chain carbonyl group in WT appears to be broken in T1032stop (Fig. 3C, left). Collectively, the LE regulates the location and orientation of both the cytosolic and TM domains in the E1·2Ca^2+^-ATP state, serving to stabilize an overall conformation unique to SERCA2b.

## Roles of the LE in structural regulation of E2P form of SERCA2b

To explore the regulatory role of the LE in E2P form of SERCA2b, we also determined and compared the cryo-EM structures of both SERCA2b WT and T1032stop bound to BeF_3_^-^. The LE in this form is located at a similar position to that in E1·2Ca^2+^-AMPPCP form (Fig. 2A and B), but displayed weaker and more fragmented density (Fig. S4), suggesting that the LE in the E2P state interacts with its neighboring regions less tightly than in the E1·2Ca^2+^-ATP state. As observed with the E1·2Ca^2+^-AMPPCP state, deletion of the LE in the E2-BeF_3_^-^ state generated two classes with significantly different conformations present in almost equal abundance (class1 = 43.4 %, class2 = 43.2 %). While neither class included Ca^2+^ at the Ca^2+^ binding sites, both showed significant density of bound BeF_3_^-^ and its neighboring Mg^2+^ in the cryo-EM maps (Fig. S5C and D), indicating that generation of the two different classes cannot be ascribed to imperfect or heterogeneous preparation of E2-BeF_3_^-^ form.

Class 1 of SERCA2b T1032stop retained almost the same arrangement for both the cytosolic and TM-helix domains as those in SERCA2b WT (Fig. 5A). In class 2, however, the cytosolic domains are tilted down toward the membrane with respect to those in class 1 (Fig. 5B). In addition to the cytosolic domain movement, class 2 underwent significant TM-helix relocation; compared with those of class 1, TM7, TM10 and TM11 of class 2 were shifted laterally to be further apart from the other TM helices (Fig. 5B, left inset), indicating a role of the LE in fine-tuning the TM helix arrangement in E2P state. Notably, class 2 adopts an overall conformation more superimposable onto the E2 form of SERCA1a rather than onto the E2P form (Fig. S7D). Thus, without the LE, the cytosolic domains can adopt alternative conformations in the E2P state, although phosphorylated D351 located at the interface of the P- and A-domains may be involved in regulating the relative arrangement of these two domains (Fig S3C).

**Fig. 5.**
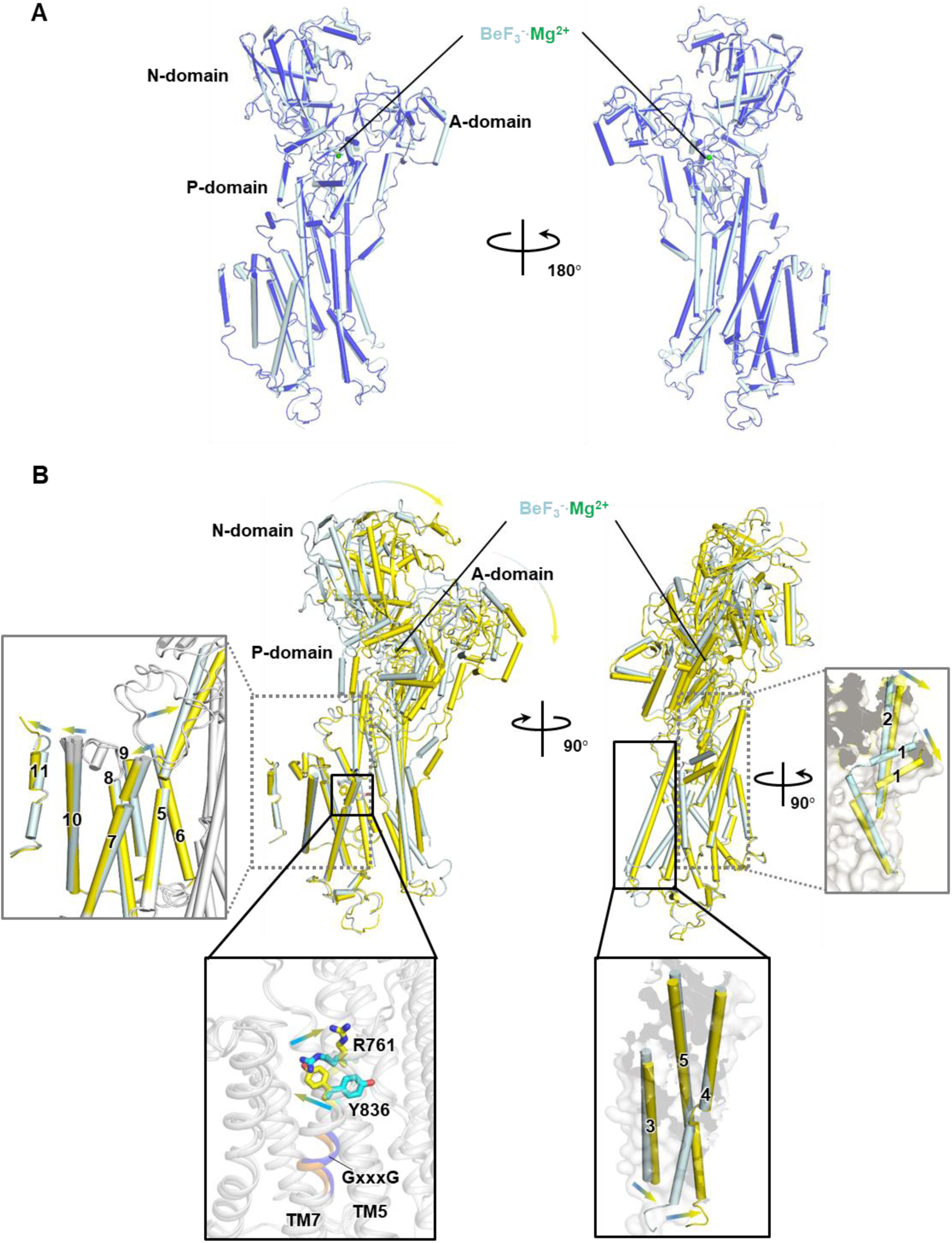
Structural effects of the LE on SERCA2b in the E2P state. **A.** Cryo-EM structures of SERCA2b WT and class 1 of T1032stop in the E2-BeF_3_^-^ state are superimposed such that the RMSD of Cα atoms in TM7-TM10 is minimized. Cylinder diagrams of SERCA2b WT and class 1 of T1032stop are colored blue and cyan, respectively. **B**. Cryo-EM structures of two different classes (class 1 and 2) of SERCA2b T1032stop in the E2-BeF_3_^-^ state are superimposed such that the RMSD of Cα atoms in TM7-TM10 is minimized. Class 1 and class 2 are colored cyan and yellow, respectively. The left upper inset presents a close-up view of TM5-TM11, in which TM helix movements during the conversion from class 1 to class 2 are indicated by arrows. The left lower inset highlights the switch residues involved in the interconversion between class 1 and class 2. The right upper and lower insets show close-up views of the movements of TM1-2 and TM3-4 during the conversion from class 1 to class 2. Note that the similar domain rearrangements take place during the transition from the E2P to E2 states in SERCA1a.

It is noteworthy that Y836 in TM7 and R761 in TM5 adopt strikingly different side-chain rotamers between class 1 and class 2 (Fig. 5B, left lower inset and Fig. S6B). Consequently, the angle of the cytosolic half of TM5 is altered (Fig. 5B, left upper inset), leading to significant inclination of the P-domain toward the membrane. In this regard, the highly conserved GxxxG motif, an element previously proposed to trigger the bending of TM5 in SERCA1a (*7, 16*), is located at the middle of TM7 in SERCA2b (Fig. 5B, left lower inset). The inclination of the P-domain induces a rotation of the A-domain due to their close contact, resulting in relocation of TM1-2 (Fig. 5B, right upper inset) and a subsequent change in orientation of TM3-4 (Fig. 5B, right lower inset). Collectively, we propose that R761 and Y836 act as switch residues that mediate conformational interconversion between class 1 and class 2 via the rotamer changes.

Importantly, a similar switching mechanism was reported to operate during the transition from E2P to E2 states in SERCA1a (*17, 18*). Hydrolysis of an aspartylphosphate at D351 allows the side-chains of R762 and Y837 to adopt different rotamers (Fig. S5E) and the subsequent rotation of the P-domain in SERCA1a, which relocates TM5 closer to TM4 (Fig. S5F). As a result, TM3 and TM4 undergo a significant positional shift to close the calcium exit channel (Fig. S5F). Concomitantly, significant reorientations of TM1 and TM2 and rotation of the A-domain are induced due to the steric effects (Fig. S5G), eventually leading to the transition to the E2 state. In contrast to SERCA2b T1032stop, the wild-type protein did not allow such conformational transitions in the E2P state, emphasizing the critical role of the LE in finely regulating the overall domain arrangement in SERCA2b.

## Proposed mechanism

Previously, TM11 of SERCA2b was reported to stabilize E2/E2P states by interacting with TM7 and TM10 (*10*). The exogenous addition of a synthetic TM11 peptide to SERCA1a resulted in a lower Vmax value and higher apparent Ca^2+^ affinity, suggesting that TM11 acts as an intramolecular uncompetitive inhibitor for SERCA2b (*19, 20*). Kinetics analysis with various SERCA2b derivatives including T1032stop suggested that whereas TM11 plays a regulatory role in the E2P dephosphorylation and the transition from E2 to E1, the LE serves to retard the E1P to E2P transition by stabilizing the E1 state via interaction with the groove formed by L5/6 and L7/8 (*20*). Thus, distinct and independent regulatory roles of TM11 and the LE were proposed.

We herein demonstrated that, despite striking discrepancy with the previous computational model (*10*), the LE is actually located outside the luminal part of TM7 and approaches a short α-helix segment of L7/8 (Fig. 1C and Fig. 2A), not the groove formed by L5/6 and L7/8, generating an overall domain arrangement that differs from SERCA1a and SERCA2a in the E1 · 2Ca^2+^-AMPPCP state. In this context, deletion of the LE rendered the overall conformation of SERCA2b similar to those of SERCA1a and SERCA2a (Fig. S7A and B), while generating multiple conformations. It is thus conceivable that the LE of SERCA2b physically fixes the E1·2Ca^2+^-ATP conformation unique to this isoform, thereby regulating the conformational transition to the E2P state (Fig. 6). Although it remains to be fully elucidated how the LE remotely modulates the orientation and location of the cytosolic domains, the overall conformation characteristic of SERCA2b appears to be stabilized most likely with specific interactions formed between R821 (L6/7) and E340 (TM5) and between R923 (L8/9) and G1013 (TM11) (Fig. 4B & C).

**Fig. 6.**
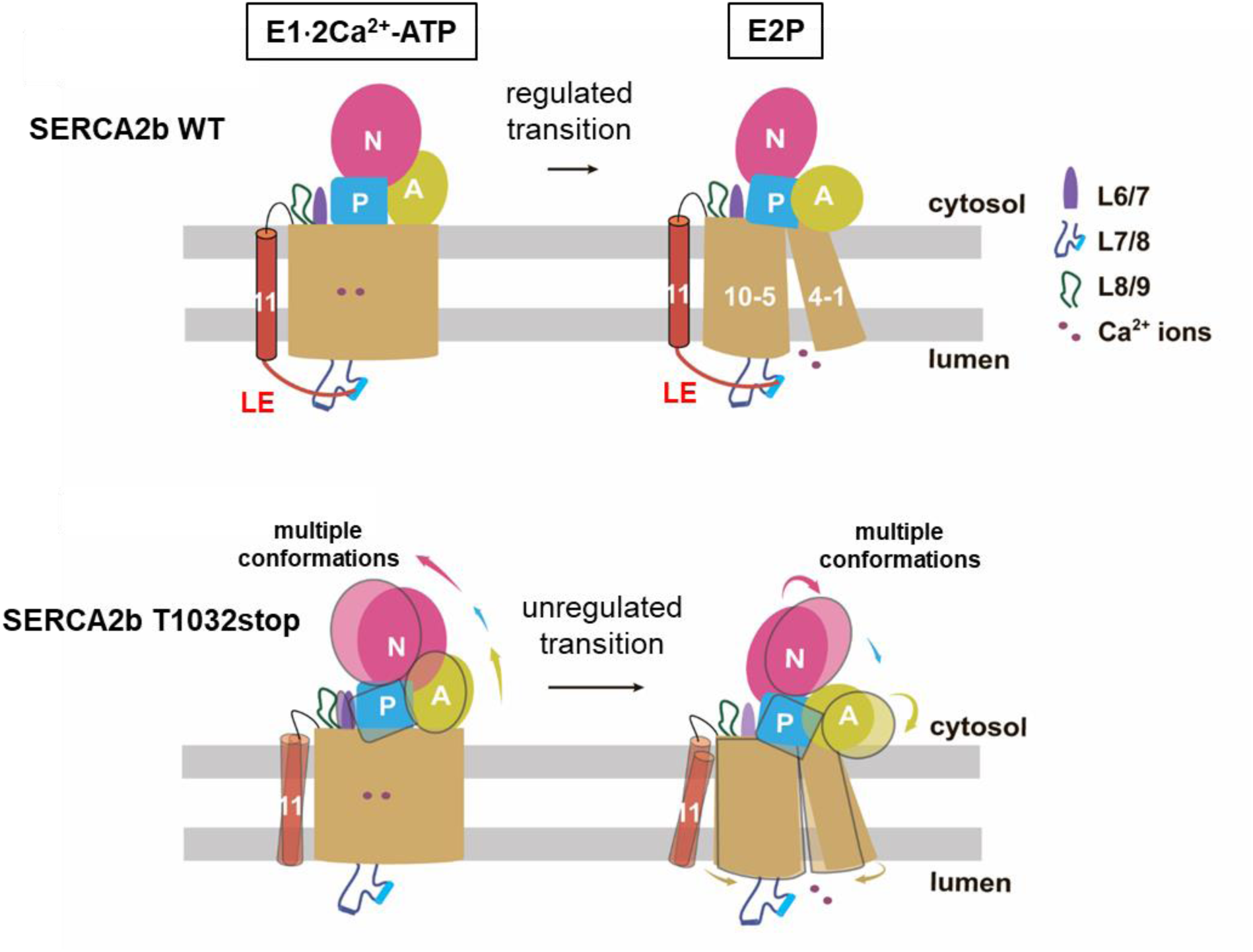
Proposed mechanisms of SERCA2b regulation by the luminal extension tail. Structures of SERCA2b WT in the E1·2Ca^2+^-ATP and E2P states are stabilized through interactions of the LE with part of L7/8 and other neighboring regions, leading to the regulated transition between these two states. Upon deletion of the LE, SERCA2b undergoes a significant relocation of both the cytosolic and TM domains accompanied by the generation of multiple overall conformations, allowing the unregulated transition from E1·2Ca^2+^-ATP to E2P states.

Alternative conformations caused by deletion of the LE were also observed in the E2P state. Although cryo-EM structures are currently unavailable for SERCA1a or SERCA2a, these isoforms also possibly adopt multiple domain arrangements due to lack of the C-terminal extension. If this is indeed the case, the LE likely plays a significant role in stabilizing the Ca^2+^-unbound form of SERCA2b, as well as the Ca^2+^-bound form, thereby modulating the kinetic properties of SERCA2b. In agreement with this notion, the rates of the dephosphorylation of E2P and the conformational transition of E2 to E1 states of SERCA2b are both considerably slower than those of SERCA1a and SERCA2a (*20*). Additionally, the overall turnover rate of SERCA2b becomes comparable to that of SERCA2a after deletion of the LE (*21*). Altogether, the regulatory role of the LE revealed in this study provides a framework for understanding the mechanism underlying the ER Ca^2+^ homeostasis ensured by the regulated cytosol-to-ER Ca^2+^ transport by SERCA2b.

## DATA AVAILABILITY

Atomic coordinates of human SERCA2b WT in E1·2Ca^2+^-AMPPCP and E2-BeF_3_^-^ states have been deposited in the Protein Data Bank under accession codes 6LLE and 6LLY, respectively. Those of three classes of human SERCA2b T1032stop in the E1·2Ca^2+^-AMPPCP state have been deposited in the Protein Data Bank under accession codes 6LN5 (class 1), 6LN6 (class 2)and 6LN7 (class 3). Those of two classes of human SERCA2b T1032stop in the E2-BeF_3_^-^ state have been deposited in the Protein Data Bank under accession codes 6LN8 (class 1) and 6LN9 (class 2). Cryo-EM density maps were deposited in the Electron Microscopy Data Bank under accession codes EMD-0912 (SERCA2b WT in E1·2Ca^2+^-AMPPCP state, EMD-0915 (SERCA2b WT in E2-BeF_3_^-^ state), EMD-0924 (class 1 of SERCA2b T1032stop in E1·2Ca^2+^-AMPPCP state), EMD-0925 (class 2 of SERCA2b T1032stop in E1·2Ca^2+^-AMPPCP state), EMD-0926 (class 3 of SERCA2b T1032stop in E1·2Ca^2+^-AMPPCP state), EMD-0927 (class 1 of SERCA2b T1032stop in E2-BeF_3_^-^ state) and EMD-0928 (class 2 of SERCA2b T1032stop in E2-BeF_3_^-^ state).

## Funding

This work was supported by CREST, JST (to K.I. and K.N., grant number JPMJCR13M6 and to M.K., grant number JPMJCR14M1), Grants-in-Aid for Scientific Research on Innovative Areas from MEXT to K.I. (18H03978) and the Basis for Supporting Innovative Drug Discovery and Life Science Research (BINDS) from the Japan Agency for Medical Research and Development (AMED) under Grant Number JP19am0101115 (support number 1025).

## Author contributions

Y.Z. performed almost all experiments, structure modelling and structure refinement. M.I. established a large-scale expression and purification system for recombinant human SERCA2b. A.T. performed acquisition and processing of cryo-EM image data. S.W. and K.I. assisted in structure modelling and refinement. T.N. assisted in data processing of cryo-EM images. K.I. supervised this work. Y.Z. and K.I. prepared figures and wrote the manuscript. Y.Z., A.T., S.W., T.N., K.N., M.K. and K.I. discussed the results and critically read the manuscript. All authors approved the manuscript for submission.

## Competing interests

We declare that there are no competing interests related to this work.

## Data Availability

Atomic coordinates of human SERCA2b WT in E1·2Ca^2+^-AMPPCP and E2-BeF_3_^-^ states have been deposited in the Protein Data Bank under accession codes 6LLE and 6LLY, respectively. Those of three classes of human SERCA2b T1032stop in the E1 · 2Ca^2+^-AMPPCP state have been deposited in the Protein Data Bank under accession codes 6LN5 (class 1), 6LN6 (class 2)and 6LN7 (class 3). Those of two classes of human SERCA2b T1032stop in the E2-BeF_3_^-^ state have been deposited in the Protein Data Bank under accession codes 6LN8 (class 1) and 6LN9 (class 2). Cryo-EM density maps were deposited in the Electron Microscopy Data Bank under accession codes EMD-0912 (SERCA2b WT in E1·2Ca^2+^-AMPPCP state, EMD-0915 (SERCA2b WT in E2-BeF_3_^-^ state), EMD-0924 (class 1 of SERCA2b T1032stop in E1·2Ca^2+^-AMPPCP state), EMD-0925 (class 2 of SERCA2b T1032stop in E1·2Ca^2+^-AMPPCP state), EMD-0926 (class 3 of SERCA2b T1032stop in E1·2Ca^2+^-AMPPCP state), EMD-0927 (class 1 of SERCA2b T1032stop in E2-BeF_3_^-^ state) and EMD-0928 (class 2 of SERCA2b T1032stop in E2-BeF_3_^-^ state).

## Supplementary Materials

### Materials and Methods

#### Expression of SERCA2b and SERCA2bT1032stop

To construct a plasmid for overexpression of SERCA2bT1032stop in HEK293T cells, a stop codon was inserted after Thr1032 using a QuickChange site-directed mutagenesis kit with primers 5’-gatctgggtctatagctgagacactaactttag-3’ and 5’-ctaaagttagtgtctcagctatagacccagatc-3’. Plasmid pcDNA3.1/PA-SERCA2bWT constructed in a previous study (22) served as the template.

Expression of wild-type (WT) SERCA2b and SERCA2bT1032stop was carried out as previously described (22). In brief, a PiggyBac Cumate Switch-Inducible Vector harboring an ORF encoding human SERCA2b WT or T1032stop were introduced into HEK293T cells along with Super PiggyBac Transposase Expression Vector (System Bioscience, LLC, CA, USA) using polyethylenimine (Sigma) to generate a stable cell line for overexpression of each protein. Cells were grown in DMEM with 4% inactivated fetal calf serum (FCS) and incubated in a humidified incubator with 5% CO2 at 37°C. After 2 days of incubation, expression of SERCA2b WT or T1032stop was induced with 150 μg/ml cumate and 50 ng/ml phorbol 12-myristate 13-acetate (PMA). Cells were incubated at 37°C for another 48 h, and then harvested by centrifugation at 1,000 × g for 15 min.

#### Purification of SERCA2b and SERCA2bT1032stop

All purification steps were performed at 4°C. Harvested cells were lysed using a Dounce Homogenizer and solubilized in a buffer containing 50 mM HEPES-NaOH (pH 7.0), 100 mM NaCl, 20% glycerol, 1 mM CaCl2, 1 mM MgCl2, 1 mM dithiothreitol (DTT), 1 mM phenylmethylsulfonyl fluoride (PMSF), and 1/100 Protease Inhibitor Cocktail (Nacalai). After homogenization, 1% (w/v) DDM was added to solubilize the membrane fraction. After 2 h of gentle rotation at 4°C, the sample was centrifuged at 14,500 rpm for 1.5 h to remove insoluble material. The supernatant was collected and incubated overnight with anti-PA sepharose beads. Beads were washed with 10 column volumes (CV) of buffer containing 50 mM HEPES-NaOH (pH 7.0), 100 mM KCl, 20% glycerol, 1 mM CaCl2, 1 mM MgCl2, 1 mM DTT, and 0.05% (w/v) lauryl maltose neopentyl glycol (LMNG), followed by elution with the same buffer containing 0.2 mg/ml PA peptide. Eluted fractions were concentrated and then further purified by size-exclusion chromatography (SEC) on a Superose 6 10/300 GL column (GE Healthcare) at 4°C with buffer containing 50 mM HEPES-NaOH (pH 7.0), 100 mM KCl, 20% glycerol, 1 mM CaCl2, 1 mM MgCl2, 1 mM DTT, and 0.05% (w/v) LMNG. Peak fractions were pooled and concentrated to 10 mg/ml with an Amicon filter device equipped with a 100 kDa cut-off membrane. Purification of SERCA2b T1032stop was performed as described for the WT protein. Representative results for SEC and SDS-PAGE are shown in Figure S1.

#### Grid preparation for cryo-EM

To prepare the E1·2Ca^2+^-AMPPCP form, 1 mM AMPPCP was added to purified SERCA2b WT or T1032stop. After overnight incubation, a second round of SEC was performed on the same column at 4°C with 50 mM HEPES-NaOH (pH 7.0), 100 mM KCl, 0% glycerol, 1 mM CaCl2, 1 mM MgCl2, 1 mM DTT, and 0.01% (w/v) LMNG to remove glycerol and decrease the detergent concentration. Preparation of the E2-BeF_3_^-^ form was performed by treating the sample with 5 mM BeSO4, 10 mM NaF, and 1 mM EGTA overnight. All samples were concentrated to 4−8 mg/ml for cryo-EM measurement. Purified samples (2−3 μl) were applied to a freshly glow-discharged 300 mesh Quantifoil 1.2/1.3 carbon grid using a Vitrobot Mark IV instrument (FEI) with a blotting time of 3−4 s under 100% humidity at 4°C, and grids were plunge-frozen in liquid ethane.

#### Cryo-EM data collection and image processing

Prepared grids were transferred to a Titan Krios G3i microscope (Thermo Fisher Scientific), which was operated at 300 kV and equipped with a Gatan Quantum-LS Energy Filter (GIF) and a Gatan K3 Summit direct electron detector in the electron counting mode. Imaging was performed at a nominal magnification of ×105,000, corresponding to a calibrated pixel size of 0.83 Å/pixel (University of Tokyo, Japan). Each movie was recorded for 3 s and subdivided into 60 frames. The electron flux was set to 14 e-/pixel/s at the detector, resulting in an accumulated exposure of 60 e-/Å^2^ at the specimen. Data were automatically acquired by the image shift method using SerialEM software, with a defocus range of −0.8 to −1.8 μm. More than 4,000 movies were acquired for each condition grid, and the total number of images is described in Supplementary Data Table 1. For all datasets, dose-fractionated movies were subjected to beam-induced motion correction using Relion-3, and contrast transfer function (CTF) parameters were estimated using CTFFIND4.

All datasets were processed using the same workflow except for 3D classification after 3D refinement. Particles were extracted from motion-corrected micrographs with down-sampling to a pixel size of 3.32 Å/pixel. These particles were subjected to one round of 2D classification and two rounds of 3D classification. The best particles were selected, re-extracted with a pixel size of 1.245 Å/pixel, and subjected to auto-3D refinement. The resulting 3D maps and particle sets were subjected to per-particle defocus refinement, beam-tilt refinement, Bayesian polishing, second per-particle defocus refinement, and 3D refinement. The global resolution of SERCA2b WT in E1·2Ca^2+^-AMPPCP and BeF_3_^-^ states was 2.9 Å and 2.8 Å, respectively. For the SERCA2b WT E1·2Ca^2+^-AMPPCP state dataset, we performed additional no-align 3D classification using a mask covering TM11 and the LE. Particles from the best class were subjected to final 3D refinement. For SERCA2b T1032stop datasets, additional local search 3D classification was performed with masks to remove micelle density. Good classes resulting from this classification were individually subjected to 3D refinement. The global resolution for SERCA2b T1032stop E1·2Ca^2+^-AMPPCP state datasets was 2.8 Å (class 1), 2.9 Å (class 2), and 2.8 Å (class 3). The global resolution for SERCA2b T1032stop BeF_3_^-^ state datasets was 3.4 Å (class 1) and 3.1 Å (class 2). All resolutions were calculated according to the FSC = 0.143 criterion. The local resolution was estimated using Relion. Detailed processing strategies are shown in Figures S7−S10.

#### Model building, refinement, and validation

Cryo-EM structures of E1·2Ca^2+^-AMPPCP and E2-BeF_3_^-^ forms of SERCA2b WT were modeled using crystal structures of human SERCA2b WT in the AMPPCP state (PDB 5ZTF) and unpublished crystal structures of SERCA2b WT in the E2-BeF_3_^-^ state as initial models, respectively. TM11 and the LE were built manually using Coot (23) based on the cryo-EM maps. After manual modification of the models, structure refinement was carried out using ‘phenix.real_space_refine’ in PHENIX (24, 25). For the E1·2Ca^2+^-AMPPCP form, linker regions composed of residues 81, 505−506, and 993−1011, which were located at L1/2, the N-domain, and L10/11, respectively, could not be modeled due to missing density in the cryo-EM map. For the E2-BeF_3_^-^ form, linker regions composed of residues 79−84, 502−507, 879−882, and 991−1014, located at L1/2, the N-domain, L7/8, and L10/11, respectively, were not modeled due to missing density in the cryo-EM map.

Cryo-EM structures of E1·2Ca^2+^-AMPPCP and E2-BeF_3_^-^ forms of SERCA2b T1032stop were built using the corresponding WT cryo-EM structures as initial models. After truncating the LE region and improving other regions of the models using Coot, structure refinement was carried out using ‘phenix.real_space_refine’ in PHENIX (*1, 2*). For the E1·2Ca^2+^-AMPPCP form, linker regions composed of residues 993−1013 contained in L10/11 could not be modeled in any of the class 1, class 2, or class 3 structures due to missing electron density in the map. Similarly, for the E2-BeF_3_^-^ form, linker regions composed of residues 78−85, 283−287, 373−377, 504−506, 876−877, and 991−1013, and residues 503−506, 876−881, and 991−1012, could not be modeled in either the class 1 or class 2 structures. The statistics for 3D-reconstitution and model refinement are summarized in Supplementary Data Table 1. Electron density for all TM helices (TM1−TM11) and cytosolic A-, N-, and P-domains, and the model-to-map FSC curves for each structural model are shown in Figures S11 and S14.

## Supplemental Figures

**Fig. S1.**
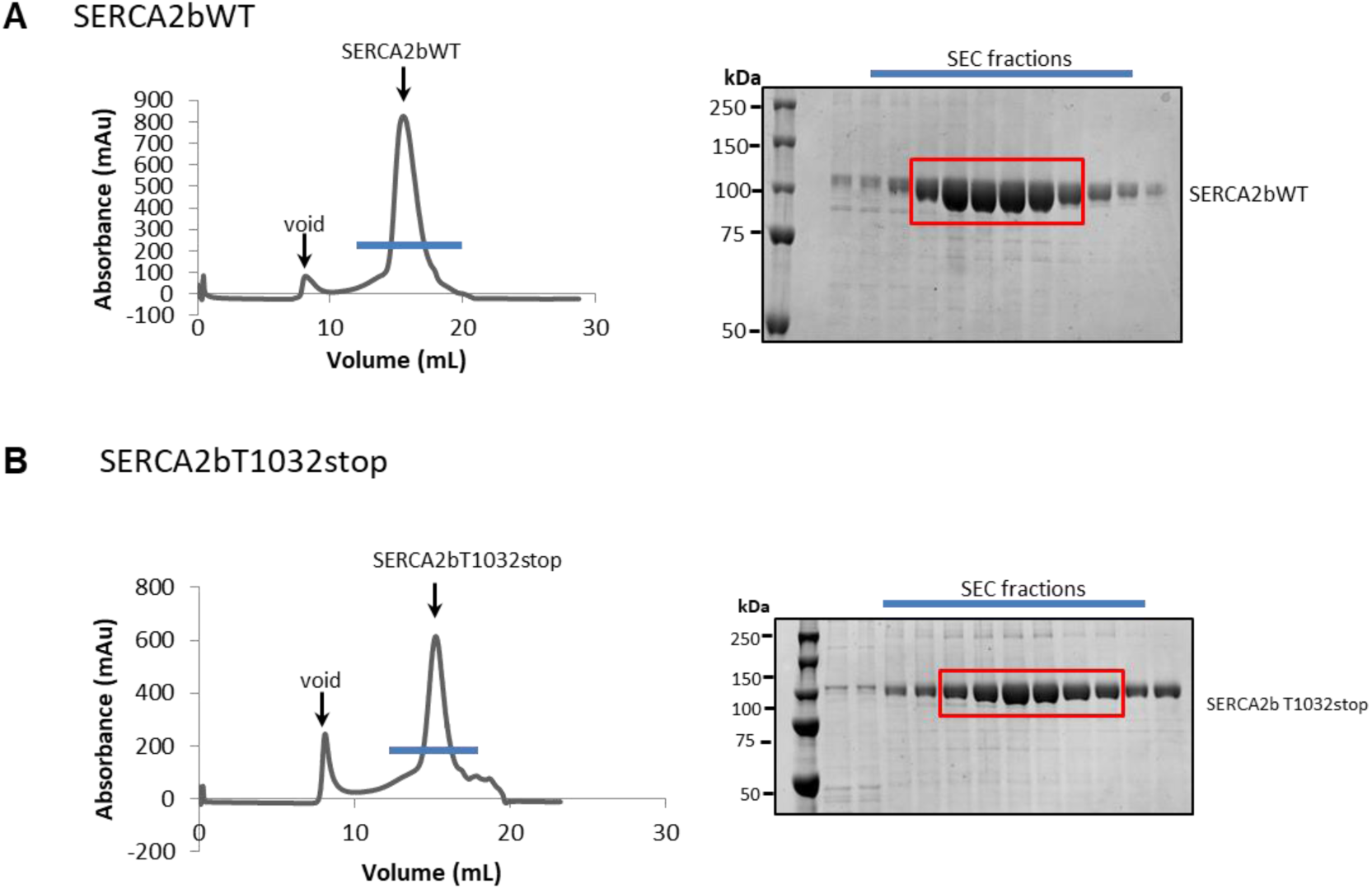
Purification of recombinant human SERCA2b. The results of SEC (left) and SDS-PAGE (right) are shown for SERCA2b WT (A) and T1032stop (B). SDS-PAGE gels were stained with Coomassie Brilliant Blue (CBB), and they show that both SERCA2b WT and T1032stop were prepared in high purity. The red boxes on gels indicate the fractions that were collected for cryo-EM measurements.

**Fig. S2.**
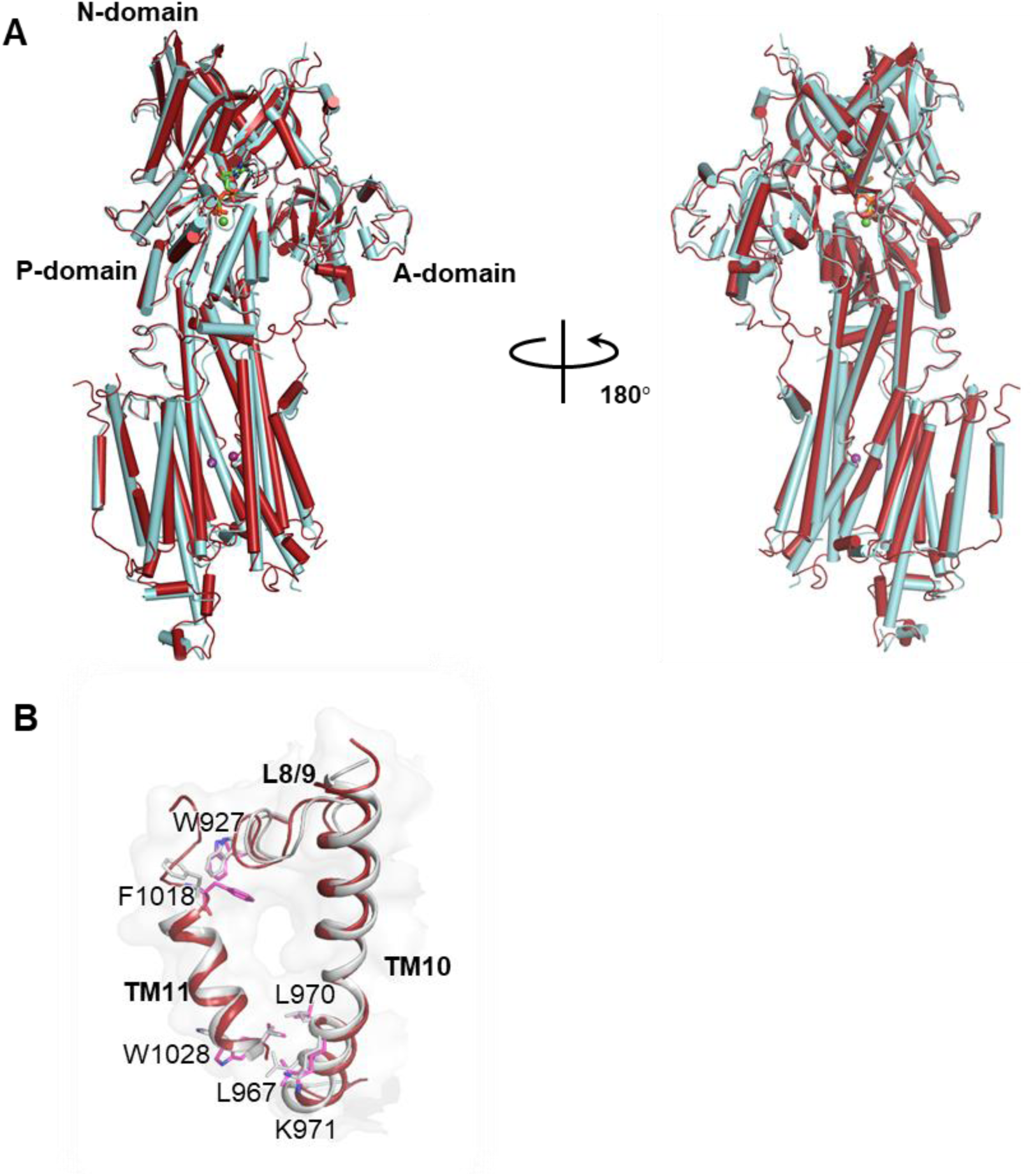
Comparison of cryo-EM and crystal structures of SERCA2b. **A.** Cryo-EM (red) and crystal (cyan; PDB ID: 5ZTF) structures of SERCA2b in the E1·2Ca^2+^-AMPPCP state are superimposed such that the RMSD of all Cα atoms is minimized. **B.** Close-up view of the residue contacts between TM11 and L8/9, and between TM11 and TM10 in the cryo-EM structure of SERCA2b in the E1·2Ca^2+^-AMPPCP state.

**Fig. S3.**
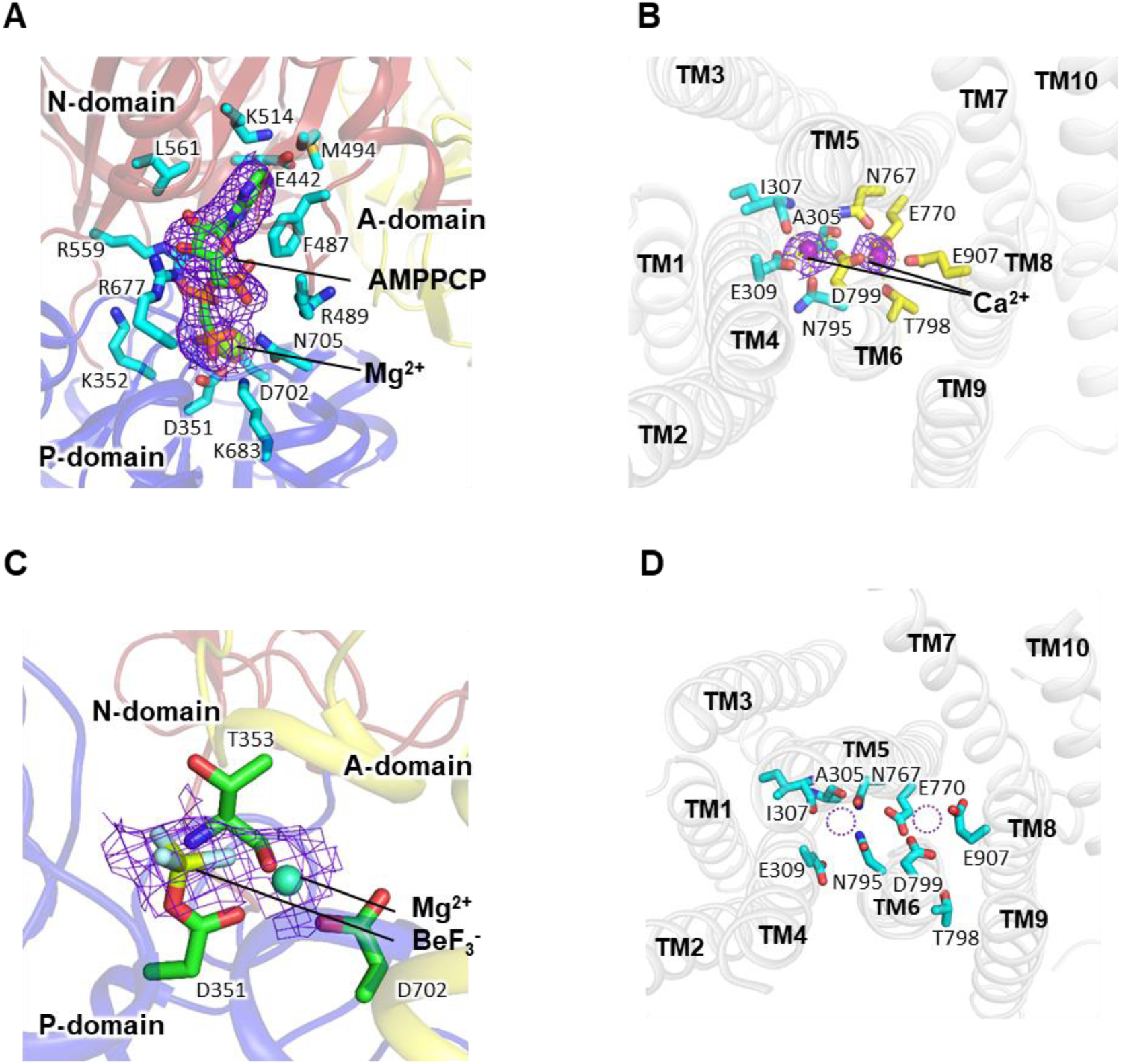
ATP- and Ca^2+^-binding sites of SERCA2b WT. **A**. Close-up view of the ATP-binding site of SERCA2b WT in the E1·2Ca^2+^-AMPPCP state, in which the bound AMPPCP molecule and its surrounding residues are represented as green and cyan sticks, respectively. The Mg^2+^ ion close to AMPPCP is depicted as a green sphere. Electron density for the bound AMPPCP is represented by violet mesh. **B.** Close-up view of the Ca^2+^-binding sites of SERCA2b WT in the E1·2Ca^2+^-AMPPCP state, in which bound Ca^2+^ ions and their electron density are represented by purple spheres and violet meshes, respectively. Neighboring residues are shown as sticks. **C.** Close-up view of the phosphorylation site of SERCA2b WT in the E2-BeF_3_^-^ state, in which the BeF_3_^-^ molecule and its electron density are represented by cyan sticks and violet mesh, respectively. Neighboring residues and the Mg^2+^ ion are represented by green sticks and spheres, respectively. **D.** Close-up view of the Ca^2+^-binding sites of SERCA2b WT in the E2-BeF_3_^-^ state. Dashed circles indicate that no Ca^2+^ is bound to SERCA2b WT in the E2P state. Neighboring residues are shown as cyan sticks.

**Fig. S4.**
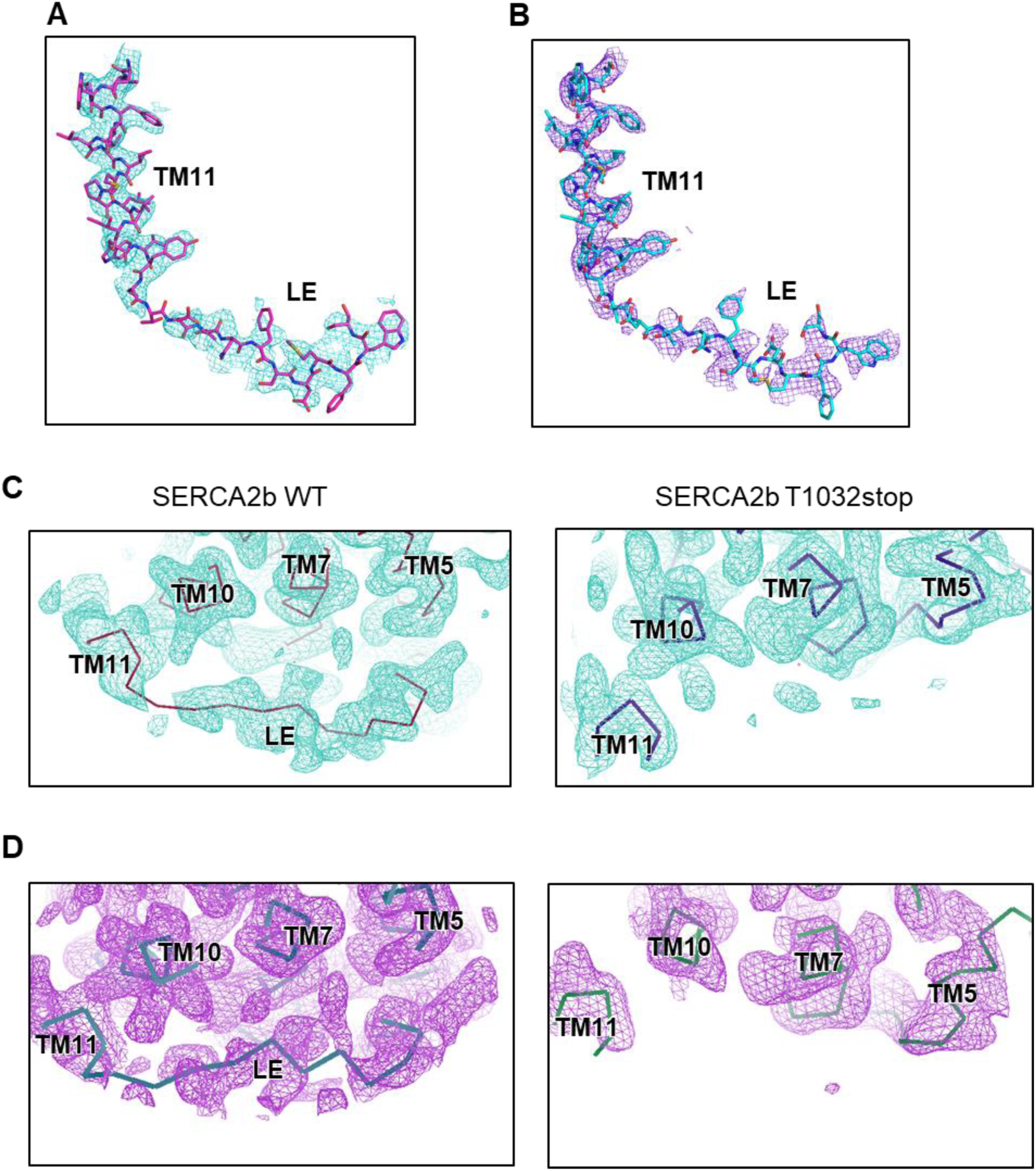
Identification of the luminal extension tail (LE) in SERCA2b. **A, B**. Cryo-EM maps at a threshold of 5.6 σ and the models of TM11 and the LE of SERCA2b WT in the E1·2Ca^2+^-AT AMPPCP (A) and E2-BeF_3_^-^ (B) states are displayed in mesh and sticks, respectively. **C.** The backbone model of the LE in SERCA2b WT in the E1·2Ca^2+^-AMPPCP state was positioned based on the cryo-EM map (left). Note that electron density is not observed around the region corresponding to the LE in the cryo-EM map of SERCA2b T1032stop (right). **D.** The backbone model of the LE in SERCA2b WT in the E2-BeF_3_^-^ state was placed based on the cryo-EM map (left). Note that electron density is not observed around the region corresponding to the LE in the cryo-EM map of SERCA2b T1032stop (right). Electron density is shown at a contour level of 6.2 RMSD in all panels.

**Fig. S5.**
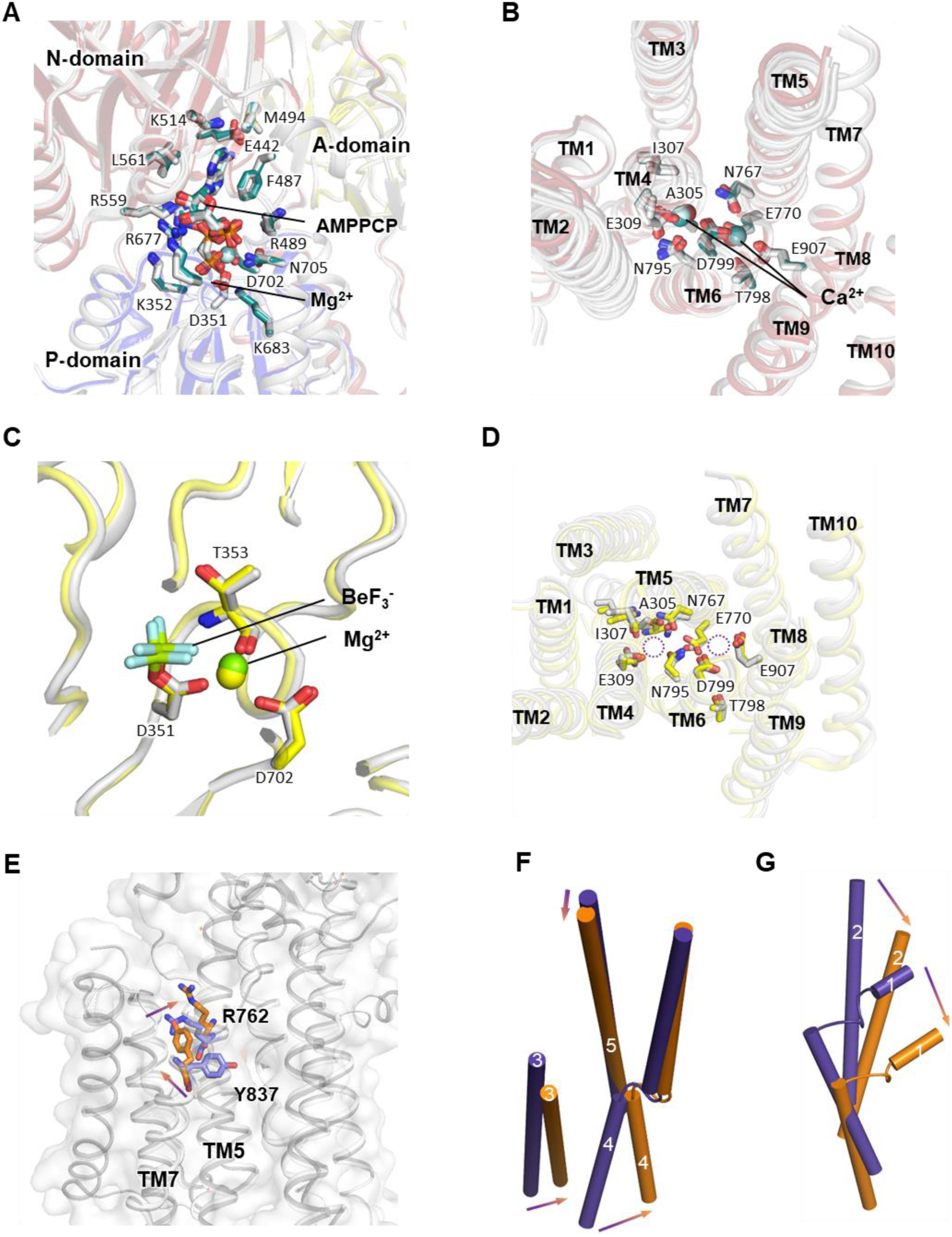
Modes of ATP, Pi, and Ca^2+^ binding in SERCA2b T1032stop. **A.** Close-up view of the ATP-binding sites of SERCA2b WT and T1032stop in the E1·2Ca^2+^-ATP state, in which the two structures are superimposed such that the RMSD of their Cα atoms in the P-domain and TM5−TM10 is minimized. The bound AMPPCP molecule and its neighboring residues are shown as cyan (WT) and gray (T1032stop) sticks. The Mg^2+^ ion close to AMPPCP is depicted as a sphere. Note that the mode of AMPPCP binding is almost identical between SERCA2b WT and T1032stop. **B.** Close-up view of the Ca^2+^-binding sites of SERCA2b WT and T1032 stop in the E1·2Ca^2+^-AMPPCP state, in which bound Ca^2+^ ions are depicted as spheres. Residues involved in Ca^2+^-binding are shown as cyan (WT) and gray (T1032stop) sticks. Note that the mode of Ca^2+^ binding is almost identical between SERCA2b WT and T1032stop. **C.** Close-up view of the phosphorylation site of class 1 and class 2 of SERCA2b T1032stop in the E2-BeF_3_^-^ state, in which the two structures are superimposed such that the RMSD of their Cα atoms in the P-domain and TM7−TM10 is minimized. The bound BeF_3_^-^ molecule is represented by cyan sticks, while its neighboring residues are represented by gray (class 1) and yellow (class 2) sticks. Note that the mode of BeF_3_^-^ binding is almost identical between class 1 and class 2. **D.** Close-up view of the Ca^2+^-binding sites of SERCA2b T1032 stop class 1 (gray) and class 2 (yellow) conformation in the E2-BeF_3_^-^ state, in which the two structures are superimposed such that the RMSD of their Cα atoms in the P-domain and TM5−TM10 is minimized. Note that no Ca^2+^ is bound at this site in either class 1 or class 2 conformations. **E.** Switch residues of SERCA1a in E2P (PDB ID: 3B9B) and E2 (PDB ID: 3W5C) states represented by purple and orange sticks, respectively. **F−G.** Movement of TM1−5 in SERCA1a during the transition from E2P to E2 states. TM helices in E2P and E2 states are colored gray and dark blue, respectively.

**Fig. S6.**
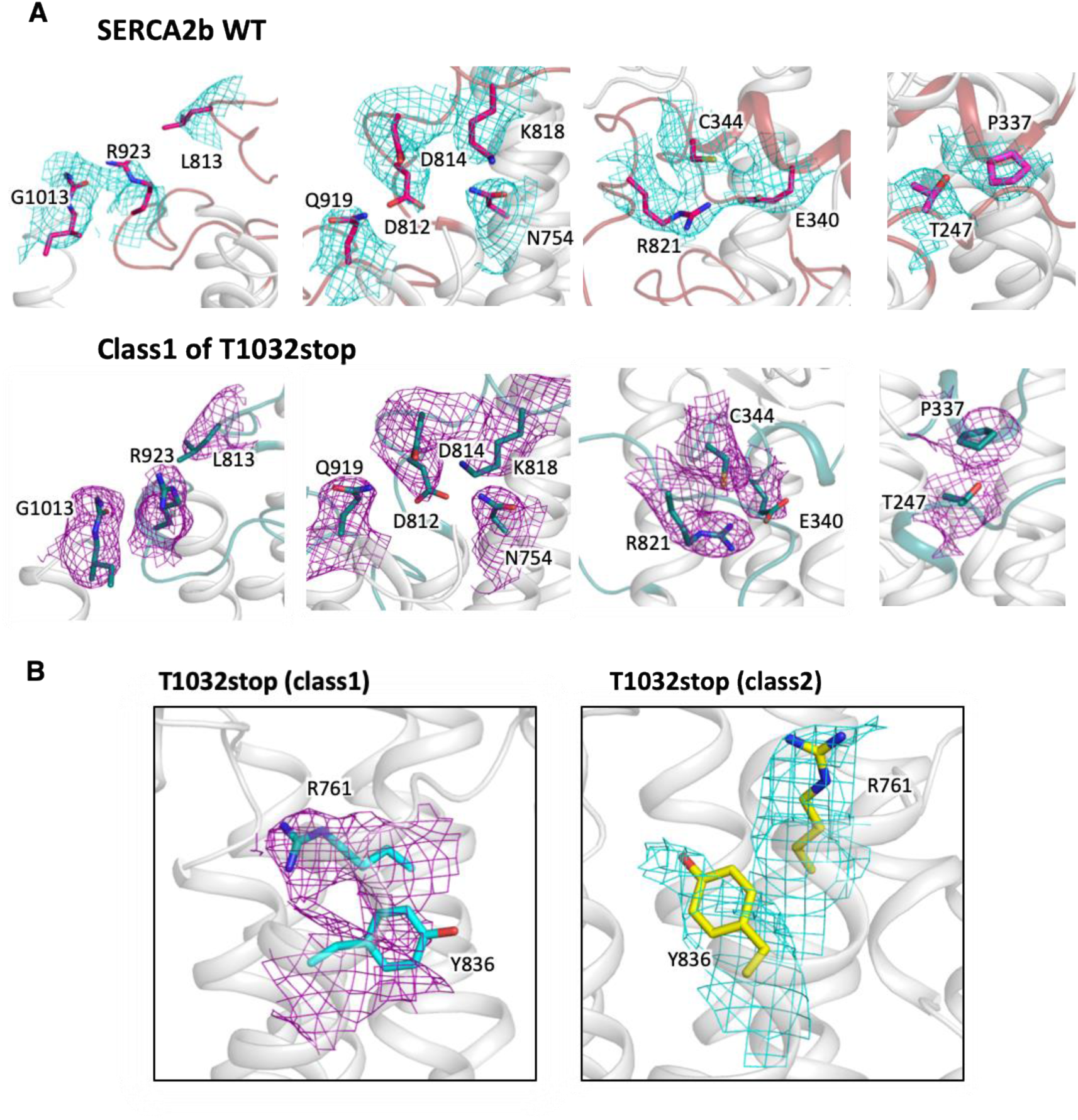
Cryo-EM density maps for critical residues involved in conformational regulation in SERCA2b WT and T1032stop. **A.** Density maps of the residues of SERCA2b in the E1·2Ca^2+^-AMPPCP state, which are focused upon in Fig. 4C, are shown at contour levels of 4.0 σ (left) and 4.5 σ (right three panels). **B.** Density maps of the switch residues in class1 (left) and class2 (right) of SERCA2b T1032stop in the E2-BeF_3_^-^ state, which are focused upon in Fig. 5B, are shown at a contour level of 4.0 σ.

**Fig. S7.**
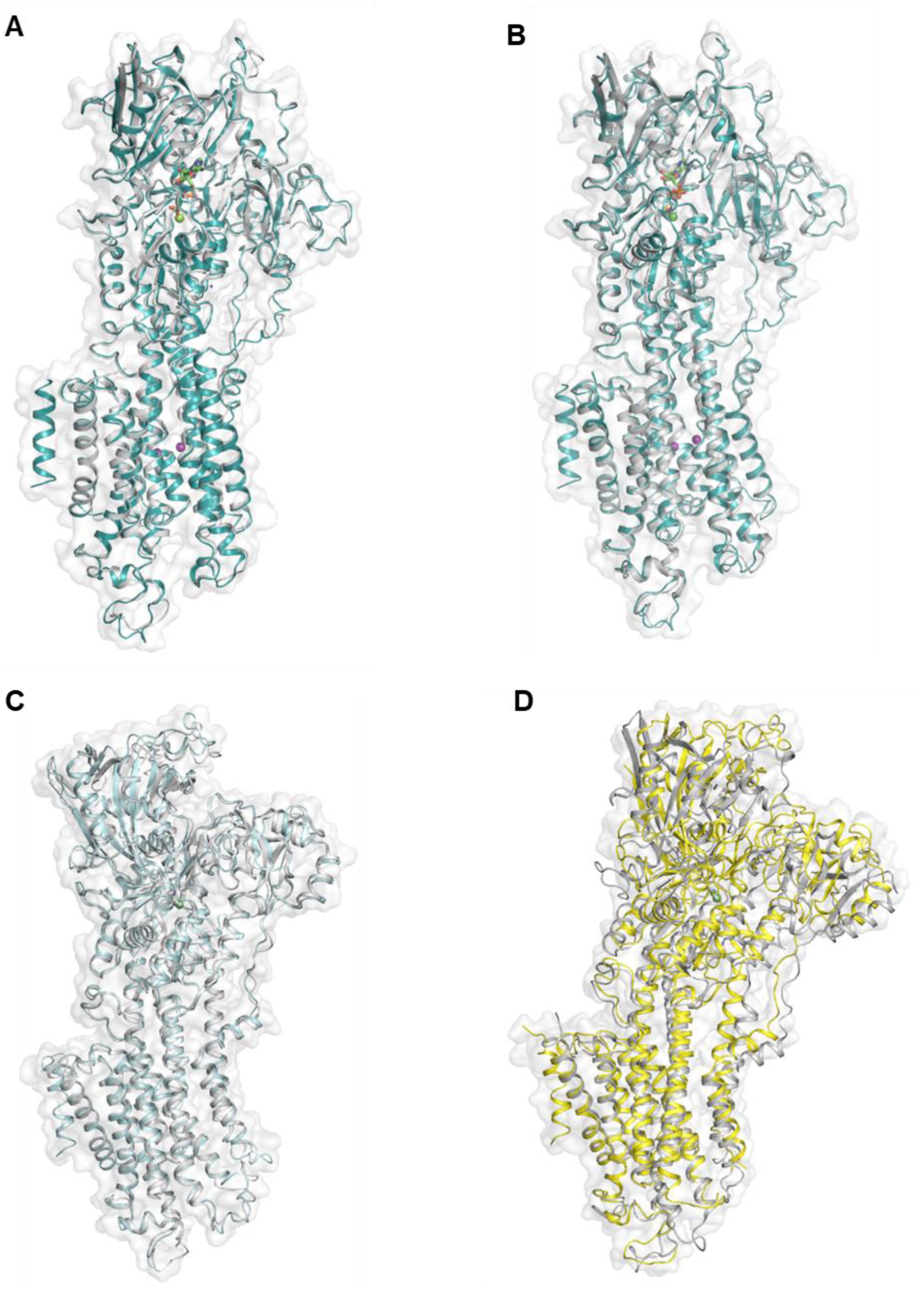
Structural similarity between SERCA2b T1032 stop and SERCA1a/SERCA2a. **A.** Alignment between the cryo-EM structure of SERCA2b T1032stop in the E1·2Ca^2+^-AMPPCP state (cyan) and the crystal structure of SERCA1a in the E1·2Ca^2+^-AMPPCP state (gray; PDB ID: 1T5S). **B.** Alignment between the cryo-EM structure of SERCA2b T1032stop in the E1·2Ca^2+^-AMPPCP state (cyan) and the crystal structure of SERCA2a in the E1·2Ca^2+^-AMPPCP state (gray; PDB ID: 6JJU). **C.** Alignment between cryo-EM structures of the class 1 conformation of SERCA2b T1032stop in the E2-BeF_3_^-^ state (cyan) and the crystal structure of SERCA1a in the E2-BeF_3_^-^ state (gray; PDB ID: 3B9B). **D.** Alignment between the cryo-EM structure of the class 2 conformation of SERCA2b T1032stop in the E2-BeF_3_^-^ state (yellow) and the crystal structure of SERCA1a in the E2 state (gray; 3W5C). All alignments were performed by superimposing the TM7−TM10 regions of the two structures of interest such that the RMSD of their Cα atoms is minimized.

**Fig. S8.**
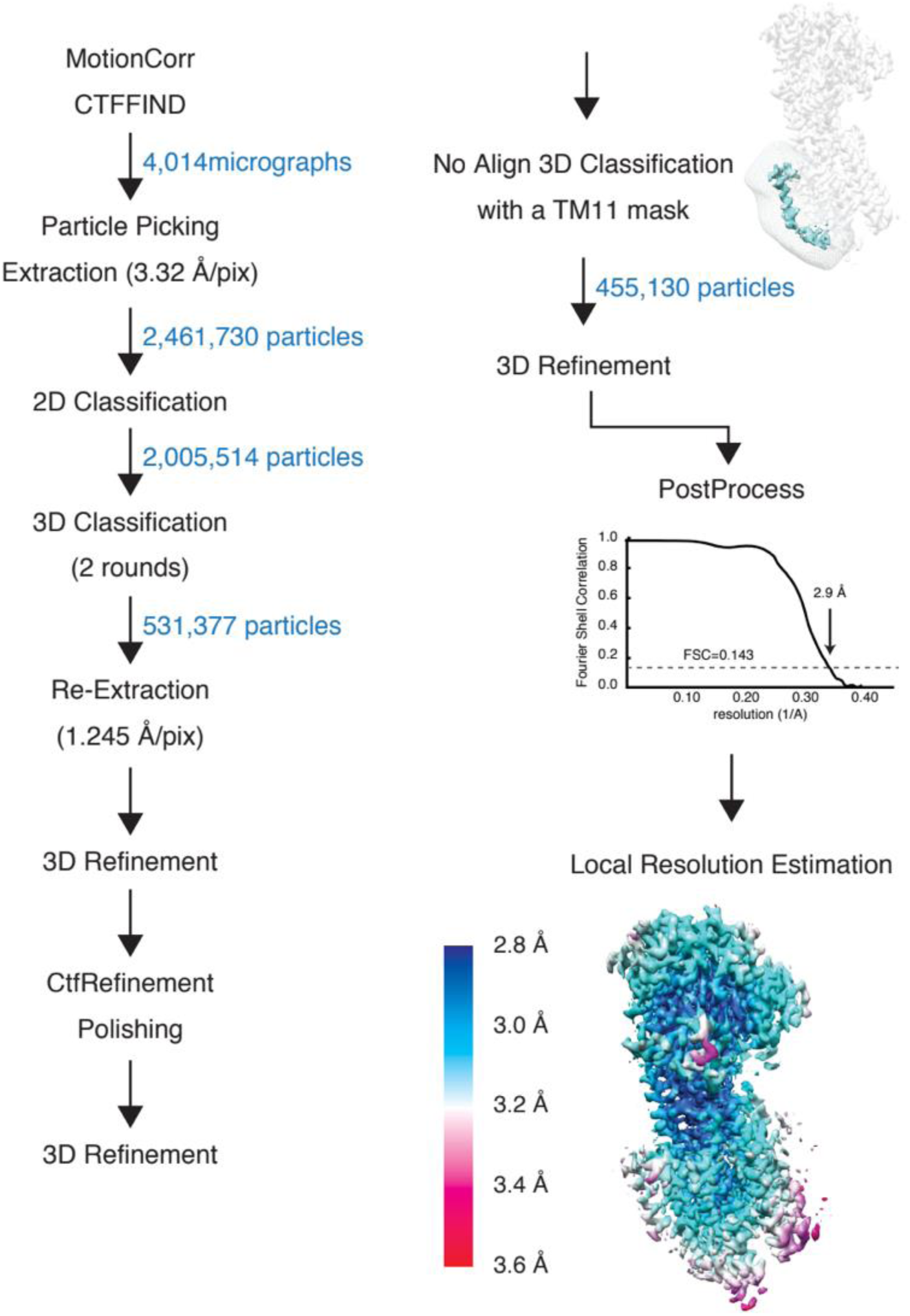
Data processing for SERCA2b WT in the E1·2Ca^2+^-AMPPCP state. The data processing workflows for single-particle image analysis and local resolution analysis is shown. Particle images were subjected to no-align 3D classification, with a mask covering TM11 and the LE. The applied mask is shown in translucent white/gray.

**Fig. S9.**
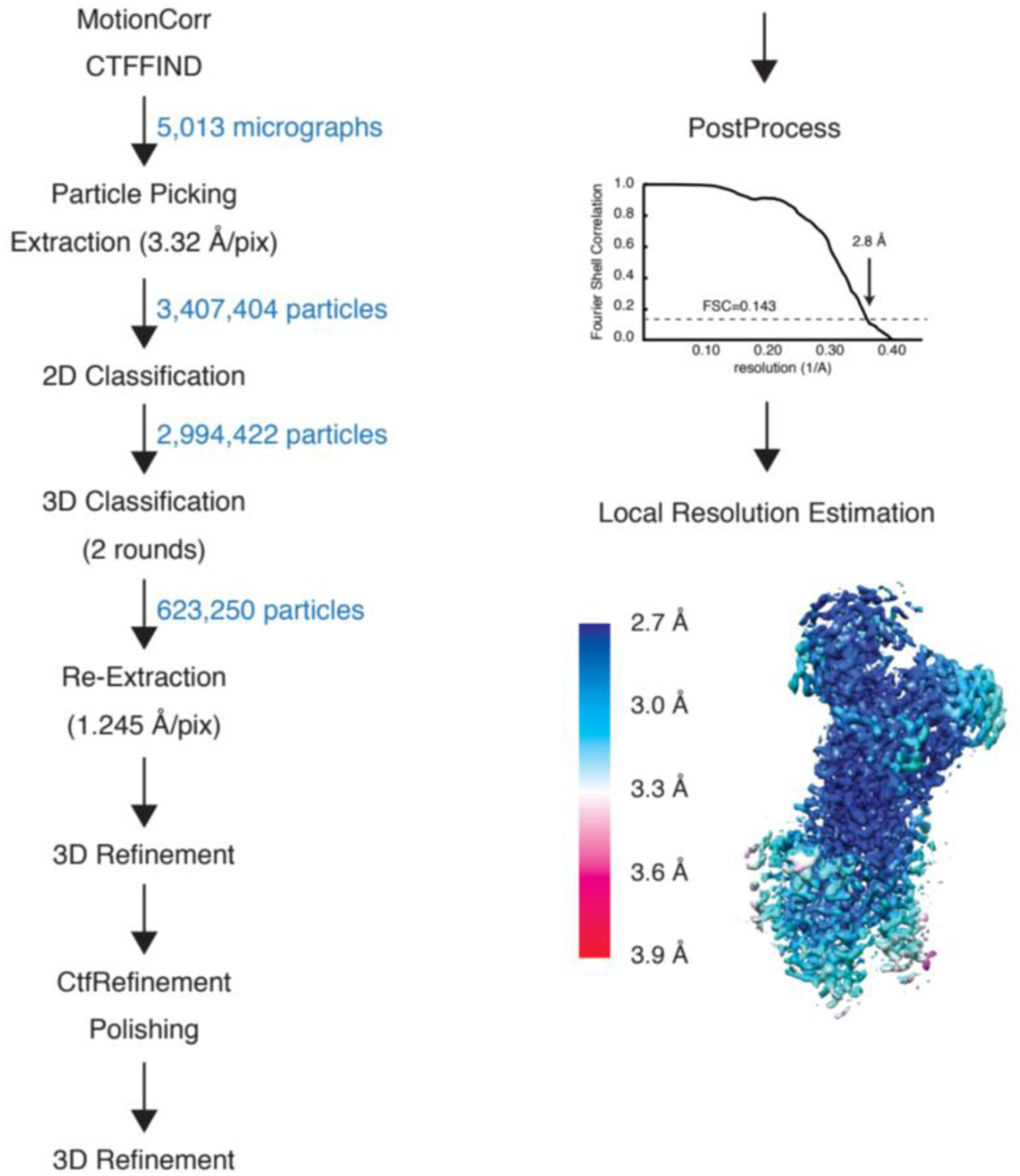
Data processing for SERCA2b WT in the E2-BeF_3_^-^ state. The data processing workflow for single-particle image analysis and local resolution analysis is shown.

**Fig. S10.**
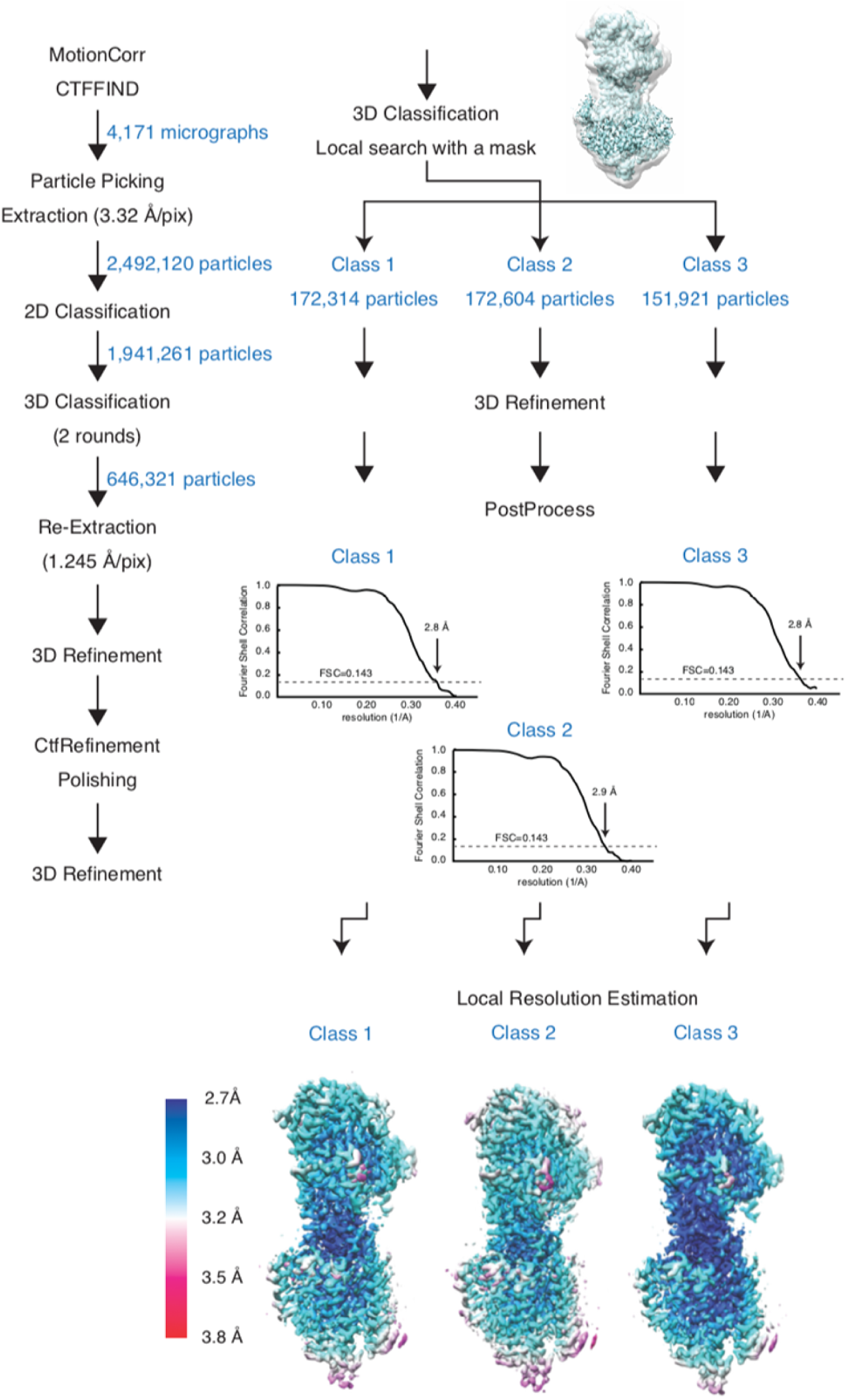
Data processing for SERCA2b T1032stop in the E1·2Ca^2+^-AMPPCP state. The data processing workflow for single-particle image analysis and local resolution analysis is shown. Particle images were separated into three groups by local search 3D classification with masks to remove micelle density.

**Fig. S11.**
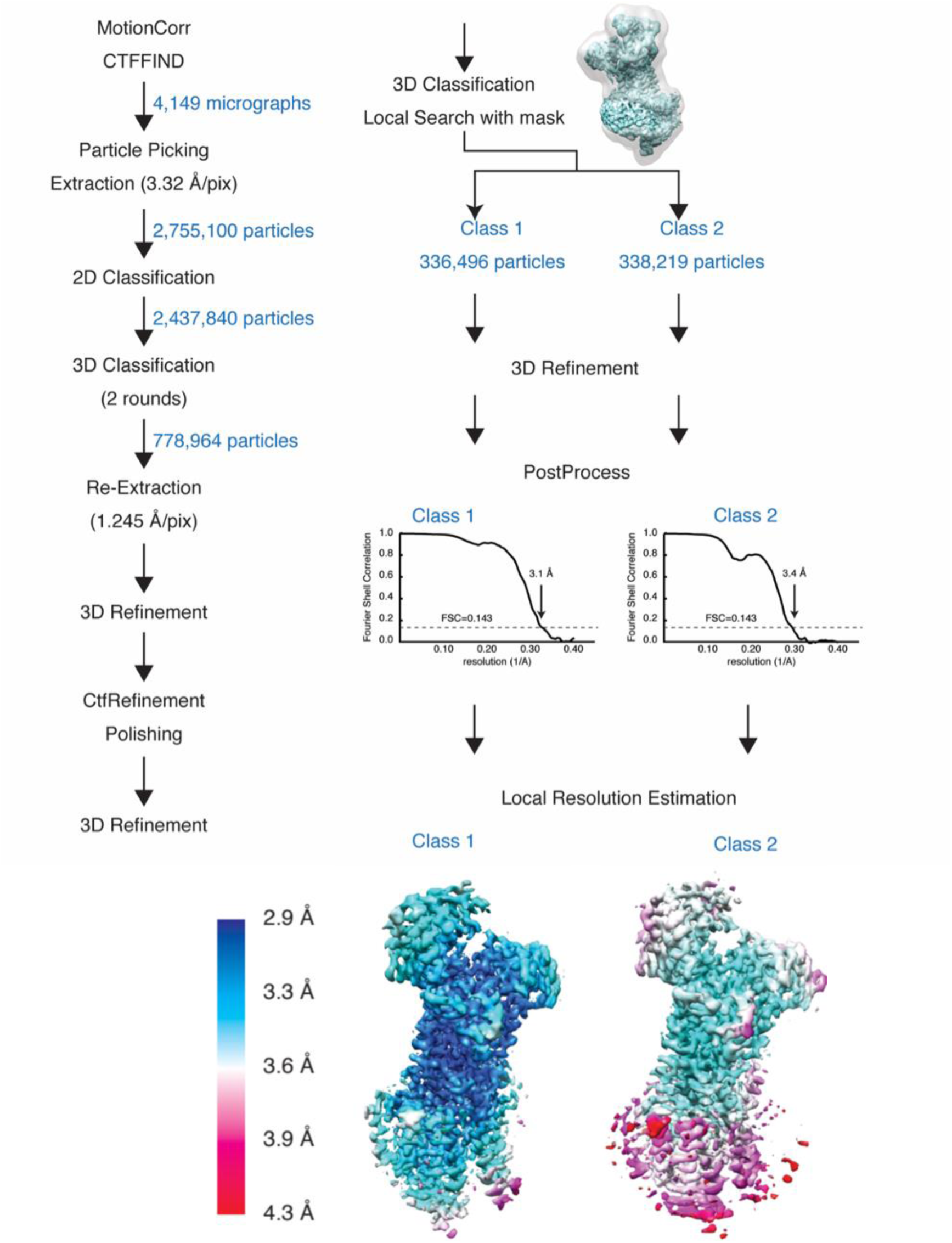
Data processing for SERCA2b T1032stop in the E2-BeF_3_^-^ state. The data processing workflow for single-particle image analysis and local resolution analysis is shown. Particle images were separated into two groups by local search 3D classification with masks to remove micelle density.

**Fig. S12.**
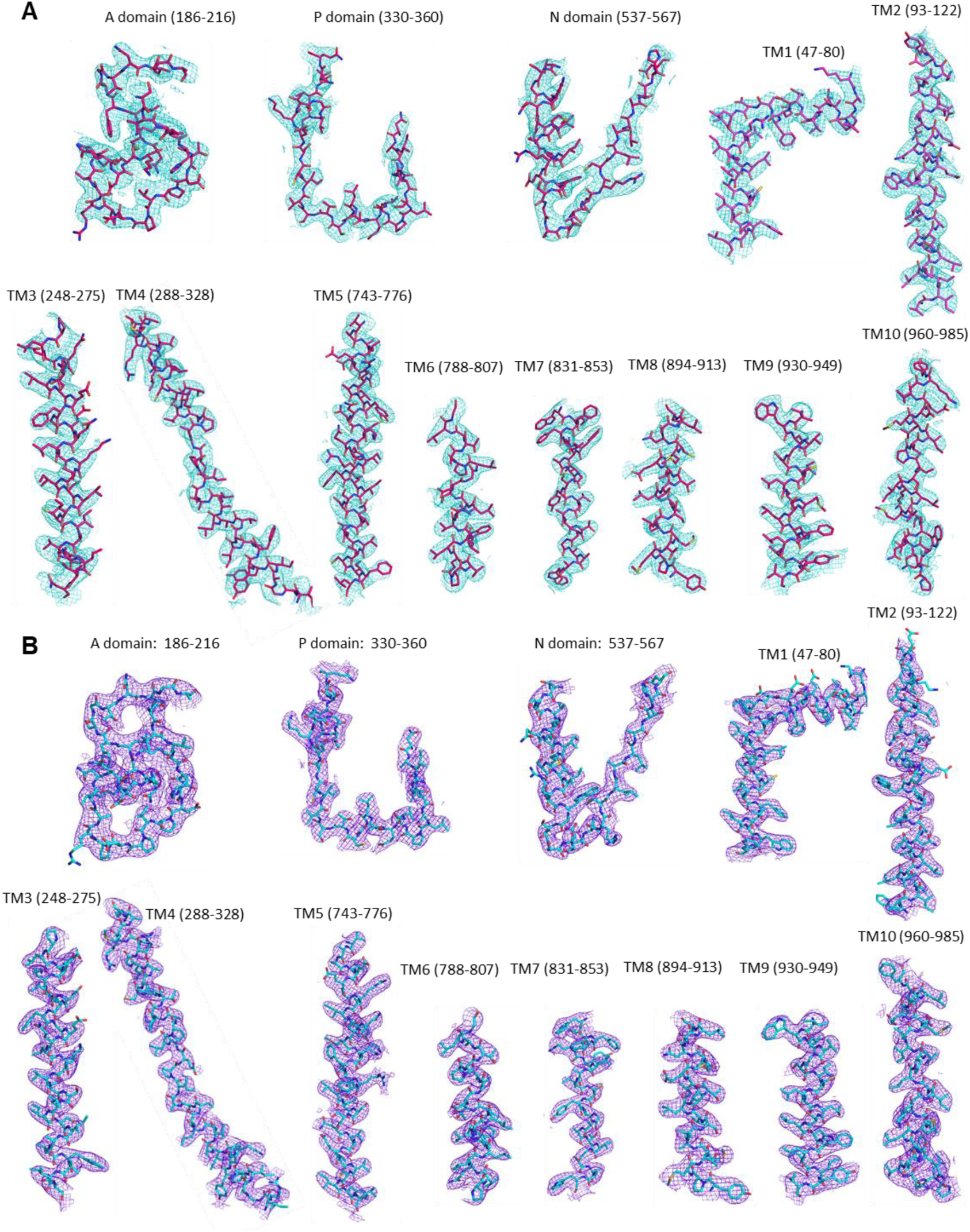
Cryo-EM density maps for TM1−11 and cytosolic A-, N-, and P-domains of SERCA2b WT in the E1·2Ca^2+^-AMPPCP (A) and E2-BeF_3_^-^ (B) states. Density is shown at a contour level of 5.6 σ for all TM helices and domains.

**Fig. S13.**
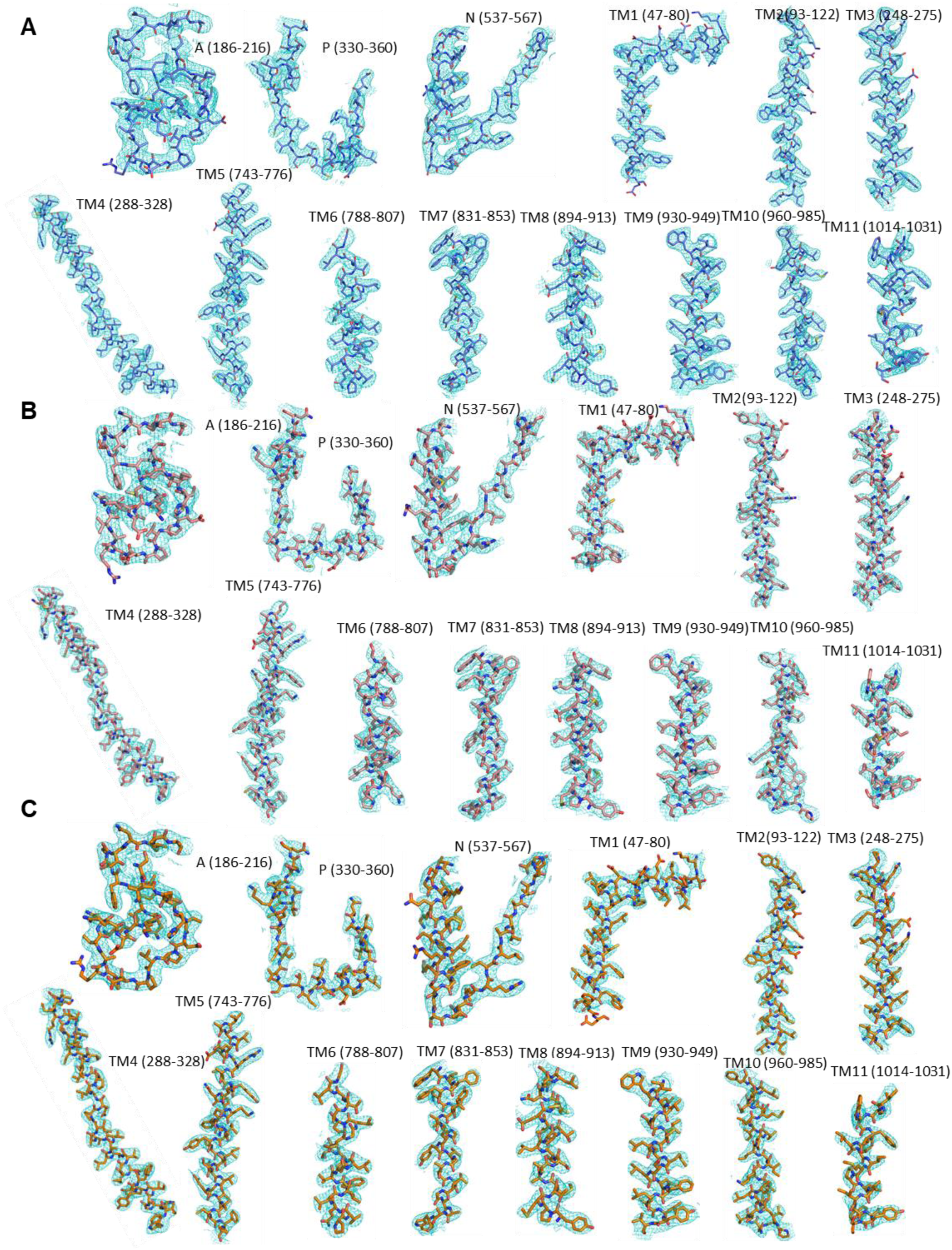
Cryo-EM density maps for TM1−11 and cytosolic A-, N-, and P-domains of SERCA2b T1032 stop in the E1·2Ca^2+^-AMPPCP state for class 1 (A), class 2 (B), and class 3 (C) conformations. Density is shown at a contour level of 5.6 σ for all TM helices and domains.

**Fig. S14.**
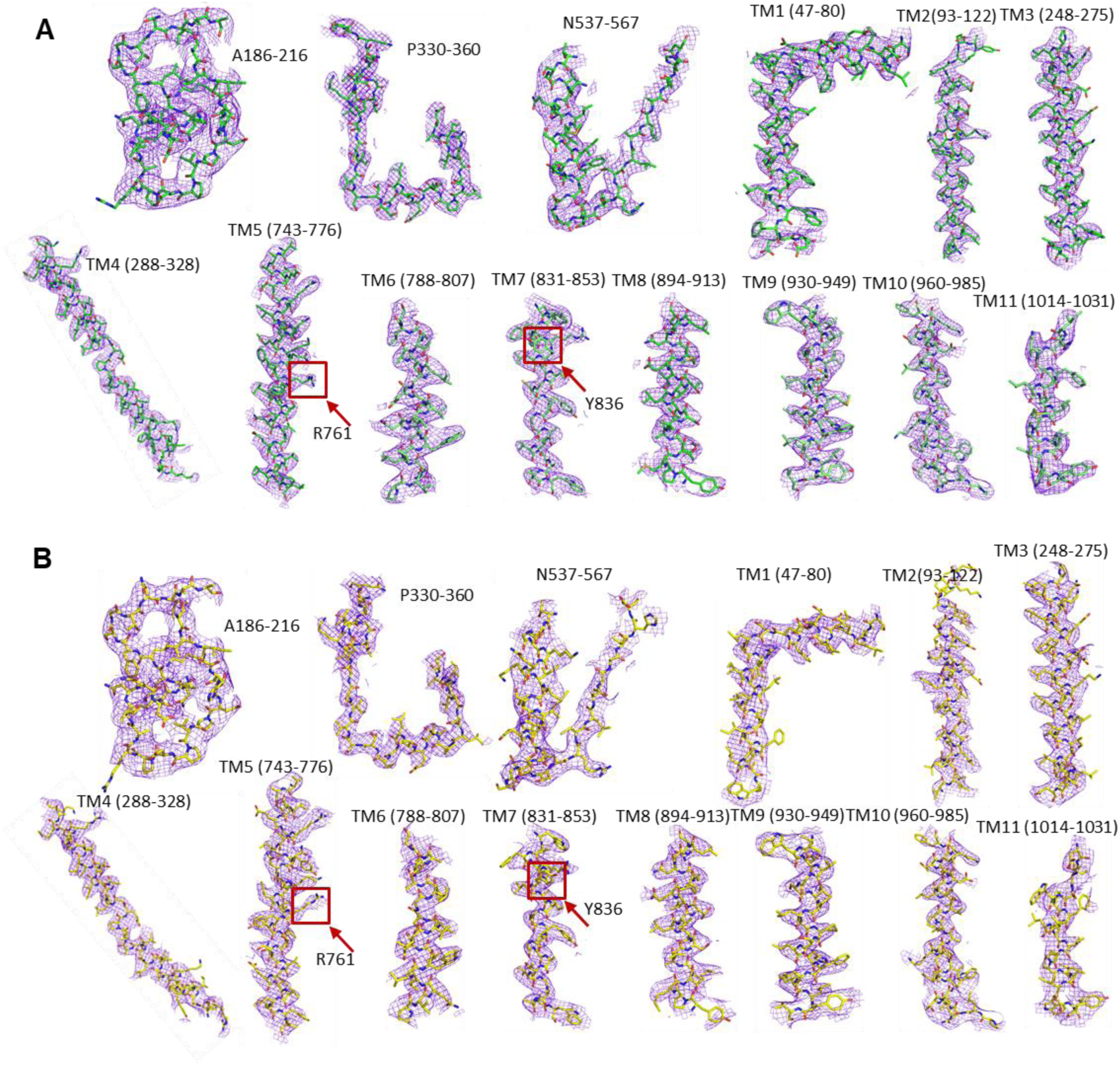
Cryo-EM density maps for TM1−11 and cytosolic A-, N-, and P-domains of SERCA2b T1032 stop in the E2-BeF_3_^-^ state for class 1 (A) and class 2 (B) conformations. Density is shown at a contour level of 5.6 σ for all TM helices and domains.

**Fig. S15.**
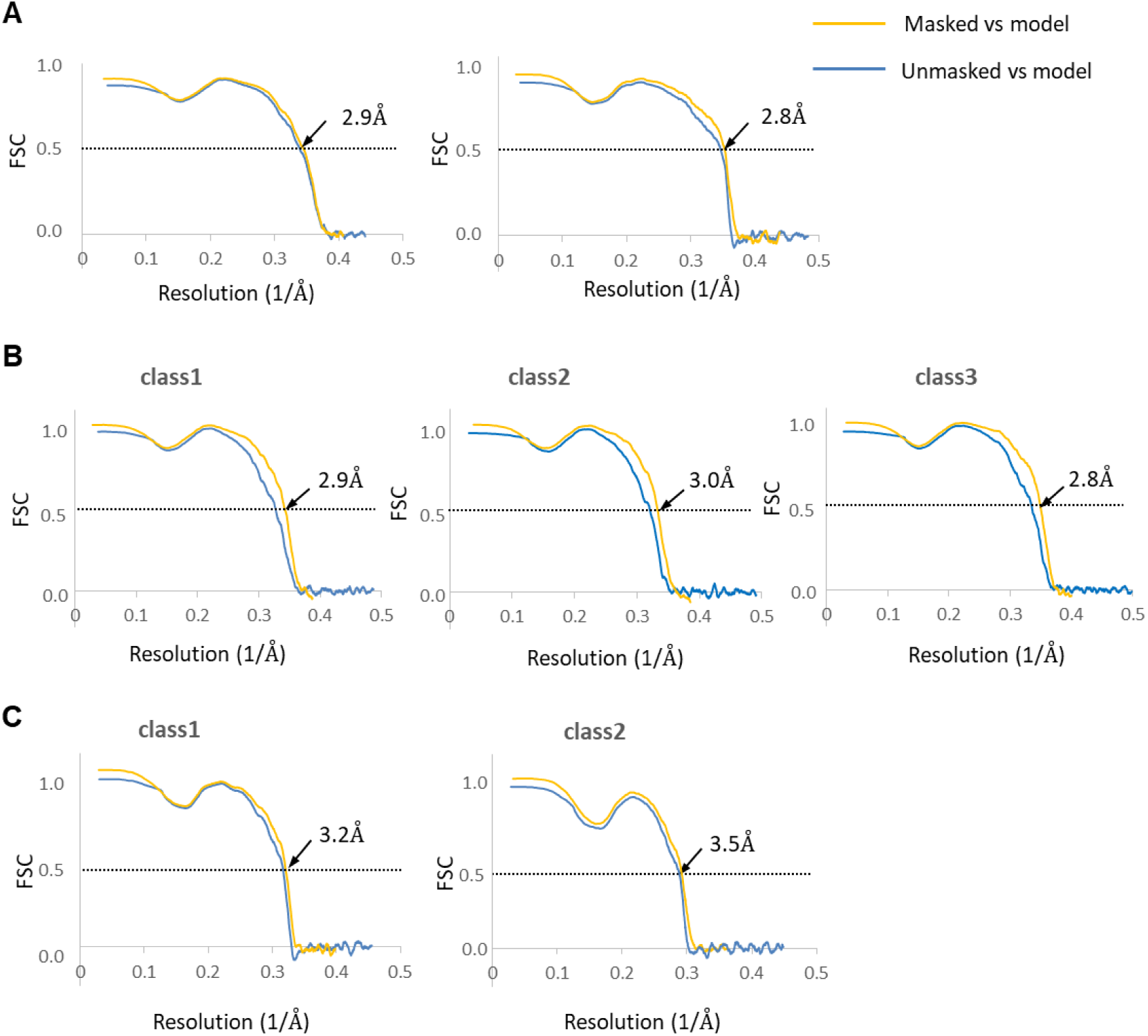
Cross-validation FSC curves for map-to-model fitting of SERCA2b WT in E1·2Ca^2+^-AMPPCP state (A) and E2-BeF_3_^-^ state (B). Cross-validation FSC curves for map-to-model fitting of T1032stop in E1**·**2Ca^2+^-AMPPCP state (C) and E2-BeF_3_^-^ state (D).

---

**Table S1.** Data collection, processing, model refinement and validation

|  | SERCA2b WT |  | SERCA2b T1032stop |  |  |  |  |
| --- | --- | --- | --- | --- | --- | --- | --- |
|  | AMP-PCP | BeF<sub>3</sub><sup>-</sup> | AMP-PCP |  |  | BeF<sub>3</sub><sup>-</sup> |  |
|  |  |  | Class 1 | Class 2 | Class 3 | Class 1 | Class 2 |
| Data collection and processing |  |  |  |  |  |  |  |
| Magnification | 105,000 |  |  |  |  |  |  |
| Voltage (kV) | 300 |  |  |  |  |  |  |
| Microscope | Titan krios G3i |  |  |  |  |  |  |
| Detector | K3 Summit |  |  |  |  |  |  |
| Electron exposure (e-/ Å) | 60 |  |  |  |  |  |  |
| Defocus range ( μ m) | -0.8 to -1.8 |  |  |  |  |  |  |
| Pixel size (Å) | 0.83 |  |  |  |  |  |  |
| Symmetry imposed | C1 |  |  |  |  |  |  |
| Number of movies | 4,014 | 5,013 | 4,171 |  |  | 4,149 |  |
| Initial particle images (no.) | 2,461,730 | 3,407,404 | 2,492,120 |  |  | 2,755,100 |  |
| Final particle images (no.) | 455,130 | 623,250 | 172,314 | 172,604 | 151,921 | 338,219 | 336,496 |
| Map resolution (Å) | 2.9 | 2.8 | 2.8 | 2.9 | 2.8 | 3.1 | 3.4 |
| FSC threshold | 0.143 | 0.143 | 0.143 | 0.143 | 0.143 | 0.143 | 0.143 |
| Map resolution range (Å) | 2.84 - 3.62 | 2.68 – 3.88 | 2.76–3.55 | 2.82 – 4.00 | 2.69 – 3.76 | 2.94 – 4.82 | 3.27 – 5.80 |
| Map sharpening B factor (Å<sup>2</sup>) | -80.5 | -58.4 | -43.2 | -63.0 | -32.0 | -66.2 | -85.9 |
| Model building and refinement |  |  |  |  |  |  |  |
| Model resolution (Å) | 2.9 | 2.8 | 2.9 | 3.0 | 2.8 | 3.2 | 3.5 |
| FSC threshold | 0.5 | 0.5 | 0.5 | 0.5 | 0.5 | 0.5 | 0.5 |
| Model Composition |  |  |  |  |  |  |  |
| Atoms | 7916 | 7774 | 7840 | 7836 | 7840 | 7739 | 7612 |
| Protein residues | 1020 | 1006 | 1011 | 1010 | 1011 | 1003 | 987 |
| Ligands | ACP:1<br>MG: 1<br>CA: 2 | BEF: 1<br>MG: 1 | ACP:1<br>MG: 1<br>CA: 2 | ACP:1<br>MG: 1<br>CA: 2 | ACP:1<br>MG: 1<br>CA: 2 | MG: 1<br>BEF: 1 | MG: 1<br>BEF: 1 |
| R.m.s. deviations |  |  |  |  |  |  |  |
| Bond length (Å) | 0.003 (0) | 0.003 (0) | 0.003 (0) | 0.003 (0) | 0.003 (0) | 0.003 (0) | 0.002 (0) |
| Bond angles (°) | 0.518 (0) | 0.565 (0) | 0.511 (0) | 0.523 (0) | 0.530 (0) | 0.521 (0) | 0.498 (0) |
| Validation |  |  |  |  |  |  |  |
| Molprobity score | 1.51 | 1.53 | 1.34 | 1.41 | 1.42 | 1.55 | 1.71 |
| Clashscore | 6.98 | 6.26 | 6.21 | 7.48 | 6.53 | 6.74 | 7.76 |
| Rotamer outliers (%) | 0 | 0 | 0 | 0 | 0 | 0 | 0 |
| EMRinger | 3.28 | 2.62 | 3.56 | 3.55 | 3.69 | 2.38 | 1.41 |
| Ramachandran Plot |  |  |  |  |  |  |  |
| Favored (%) | 97.34 | 96.89 | 98.01 | 98.01 | 97.72 | 96.99 | 95.89 |
| Allowed (%) | 2.66 | 3.11 | 1.99 | 1.99 | 2.28 | 3.01 | 4.11 |
| Outliers (%) | 0 | 0 | 0 | 0 | 0 | 0 | 0 |
| CaBLAM outliers (%) | 0.99 | 0.81 | 0.20 | 0.70 | 0.9 | 0.91 | 0.52 |

